# Nitric oxide coordinates histone acetylation and expression of genes involved in growth/development and stress response

**DOI:** 10.1101/2020.06.16.154476

**Authors:** Alexandra Ageeva-Kieferle, Elisabeth Georgii, Barbro Winkler, Andrea Ghirardo, Andreas Albert, Patrick Hüther, Alexander Mengel, Claude Becker, Jörg-Peter Schnitzler, Jörg Durner, Christian Lindermayr

**Author notes:** Lead Contact: PD Dr. Christian Lindermayr, Institute of Biochemical Plant Pathology, Helmholtz Zentrum München, Ingolstädter Landstrasse 1, 85764 München/Neuherberg.

## Abstract

Nitric oxide (NO) is a signaling molecule with multiple regulatory functions in plant physiology and stress response. Besides direct effects on the transcriptional machinery, NO can fulfill its signaling function via epigenetic mechanisms.

We report that light intensity-dependent changes in NO correlate with changes in global histone acetylation (H3, H3K9 and H3K9/K14) in *Arabidopsis thaliana* wild-type leaves and that this correlation depends on S-nitrosoglutathione reductase and histone deacetylase 6. The activity of histone deacetylase 6 was sensitive to NO, which demonstrates that NO participates in regulation of histone acetylation. ChIP-seq and RNA-seq analyses revealed that NO is involved in the metabolic switch from growth and development to stress response. This coordinating function of NO might be of special importance in adaptation to a changing environment and could therefore be a promising starting point to mitigating the negative effects of climate change on plant productivity.

## 1. Introduction

Nitric oxide (NO) is a ubiquitous signaling molecule with pleiotropic functions throughout the lifespan of plants. Indeed, NO is involved in several physiological processes, including growth and development, but also in iron homeostasis as well as biotic and abiotic stress response, such as to high salinity, drought, UV-B radiation, high temperature, and heavy metal toxicity (An et al., 2005; Besson-Bard et al., 2009; Delledonne et al., 1998; Durner et al., 1998; Mata and Lamattina, 2001; Puyaubert and Baudouin, 2014; Tian et al., 2007; Zhao et al., 2004; Zhao et al., 2007). NO is a heteronuclear diatomic radical with a half-lifetime of 3-5 seconds in biological systems and the multifunctional role of NO is based on its chemical properties, cellular environment, and compartmentalization. Depended to a large extent on its local concentration, which is affected by its rate of synthesis, displacement, and removal, NO has been described as cytoprotective, signaling, or cytotoxic molecule (Ageeva-Kieferle et al., 2019; Buet and Simontacchi, 2015; Fancy et al., 2017; Floryszak-Wieczorek et al., 2006; Mur et al., 2013; Trapet et al., 2015; Yu et al., 2014).

NO fulfills its biological functions by modulating protein function/activity through different types of post-translational modifications (PTM): Protein S-nitrosation, tyrosine nitration or metal nitrosylation. Protein S-nitrosation – the covalent attachment of NO to the sulfur group of cysteine residues – is one of the most important NO-dependent protein modifications, and plants respond to many different environmental changes by S-nitrosating a specific set of proteins (Jain et al., 2018; Puyaubert et al., 2014; Romero-Puertas et al., 2008; Vanzo et al., 2016). S-Nitrosated glutathione (S-nitrosoglutathione, GSNO) has an important function as NO reservoir, NO transporter, and physiological NO donor, which can transfer its NO moiety to protein cysteine residues (Hess et al., 2005; Kovacs and Lindermayr, 2013). Therefore, the level of S-nitrosated proteins correlates with GSNO levels. The level of GSNO is controlled by the catalytic activity of GSNO reductase (GSNOR; EC: 1.1.1.284). This enzyme is catalysing the degradation of GSNO to oxidized glutathione and ammonium and in this way regulates directly the level of GSNO and indirectly the level of S-nitrosated proteins (Liu et al., 2001; Sakomoto et al., 2002). Loss of GSNOR function results in enhanced levels of low and high molecular S-nitrosothiols (SNOs) (Feechan et al., 2005; Kovacs et al., 2016; Lee et al., 2008). The pleiotropic phenotype of *GSNOR-knock-out* mutants (background Columbia and Wassilijewskija) and their sensitivity to biotic and abiotic stress clearly demonstrate the importance of this enzyme for plant growth, development and stress response (Feechan et al., 2005; Holzmeister et al., 2011; Kwon et al., 2012; Lee et al., 2008; Wünsche et al., 2011; Xu et al., 2013).

Using a site-specific nitrosoproteomic approach, several hundred target proteins for S-nitrosation were identified in *A. thaliana gsnor* plants (Hu et al., 2015). These proteins are involved in a wide range of biological processes and amongst others play a role in chlorophyll metabolism and photosynthesis. Consistently, *gsnor* mutants showed altered photosynthetic properties, such as increased quantum efficiency of photosystem II (PSII) photochemistry and photochemical quenching, and decreased non-photochemical quenching (Hu et al., 2015), suggesting that S-nitrosation is an important regulatory mechanism for light-dependent processes. In several studies, *gsnor* plants have been analyzed on proteome and transcriptome levels to gain insights into the physiological functions of this enzyme (Fares et al., 2011; Holzmeister et al., 2011; Kovacs et al., 2016; Kuruthukulangarakoola et al., 2017).

Gene transcription can be regulated via modification of transcription factors or via chromatin modifications. The chromatin structure in eukaryotic organisms is very dynamic and changes in response to environmental stimuli. Chromatin marks are defined modifications on histone tails or DNA, playing key roles in processes such as gene transcription, replication, repair, and recombination (Bannister and Kouzarides, 2011). DNA methylation is usually associated with long-term silencing of genes, whereas histone modifications contribute to both activation and repression of gene transcription and are mostly removed after several cell cycles (Jaenisch and Bird, 2003; Minard et al., 2009). Several lines of evidence demonstrate that NO regulates gene expression via modification of the chromatin structure and/or DNA accessibility. In general, the distinct chromatin states that modulate access to DNA for transcription are regulated by multiple epigenetic mechanisms, including DNA methylation, covalent modifications of core histones such as methylation and acetylation, ATP-dependent chromatin remodeling, placement of histone variants, non-coding RNAs, and metabolo-epigenetic effects (Lindermayr et al., 2020; Schvartzman et al., 2018; Zhang et al., 2018). Recently, we demonstrated that NO affects histone acetylation by targeting and inhibiting histone deacetylase (HDA, EC: 3.5.1.98) complexes, resulting in the hyperacetylation of specific genes (Mengel et al., 2017). Treatment with the physiological NO donor GSNO increased global histone 3 (H3) and histone 4 (H4) acetylation. Chromatin immunoprecipitation sequencing (ChIP-seq) revealed that several hundred genes displayed NO-regulated histone acetylation. Many of these genes were involved in plant defense response and abiotic stress response, but also in chloroplast function, suggesting that NO might regulate expression of specific genes by modulation of chromatin structure (Mengel et al., 2017).

Arabidopsis contains 18 members of HDAs, divided into three subfamilies: RPD3-like, HD-tuins and sirtuins (Hollender and Liu, 2008). The first subfamily is the largest one and is composed of twelve putative members (HDA2, HDA5-10, HDA14-15, HDA17-19), which, based on structural similarity, can be further divided into three classes. HDAs of this type are homologous to yeast reduced potassium deficiency 3 (RPD3) proteins that are ubiquitously present in all eukaryotes. All members of this subfamily contain a specific deacetylase domain that is required for their catalytic activity. The second subfamily contains the HD-tuins (HD2) and was originally found in maize. This type of proteins is plant-specific, although homologous cis-trans prolyl isomerases are also present in other eukaryotes (Dangl et al., 2001). The third subfamily of plant HDAs is represented by sirtuins (SIR2-like proteins), which are homologs to yeast silent information regulator 2 (SIR2) (Pandey et al., 2002). These HDAs are unique because they require a NAD cofactor for functionality, and unlike RPD3 proteins, they are not inhibited by trichostatin A (TSA) or sodium butyrate. Moreover, sirtuins use a wide variety of substrates beyond histones.

Here, we report that increased light intensity (dark, 200 µmol photons m^-2^s^-1^, 1000 µmol photons m^-2^s^-1^) enhances NO/SNO levels in Arabidopsis leaves. These light intensity-dependent changes in SNO/NO levels positively correlate with changes in global H3 acetylation and acetylation of H3K9 and H3K9/K14 in Arabidopsis wild-type (wt) plants. Interestingly, there were no light intensity-dependent changes in histone acetylation observed in plants with loss of GSNOR (*gsnor1-3*, (Feechan et al., 2005; Lee et al., 2008) or HDA6 (*axe1-5*; (Murfett et al., 2001; Wu et al., 2008), a member of the RPD3-like subfamily, suggesting a light intensity-dependent regulatory function of GSNOR and HDA6 on histone acetylation. *In vitro* measurement of enzyme activities provided evidence for NO-sensitivity of Arabidopsis HDA6. A ChIP-seq analysis of the H3K9ac mark in wt, *gsnor* and *hda6* mutants under dark und low light conditions identified 16,276 acetylated loci. Interestingly, under low light, GSNOR and HDA6 share a significant function in deacetylation of genes involved in growth/development and acetylation of stress-responsive genes, suggesting a link between GSNOR (level of SNO) and HDA6 in these functions. Furthermore, RNA-seq analysis of wt, *gsnor* and *hda6* mutants under these conditions revealed a common function of GSNOR and HDA6 in downregulation of genes involved in growth/development and in the same time upregulation of stress-related genes. In summary, our data suggest a function for NO as molecular switch between growth/development on one side and stress responses on the other side.

## 2. Results

### 2.1. Enhanced light-dependent production of NO/SNO in Arabidopsis plants with loss of GSNOR function

In plants, both NO and radiation are important regulators of growth, development and stress response. To further demonstrate a link between light and NO/SNOs, NO emission of single *Arabidopsis* plants was analyzed under different light intensities in a closed cuvette using a CLD Supreme NO analyzer. Only very small differences in NO emission between low light (LL, photosynthetic photon flux density (PPFD) of 200 µmol photons m^-2^s^-1^, T 22°C) and dark (D, PPFD of 0 µmol photons m^-2^s^-1^, T 22°C) conditions were observed in Arabidopsis wt plants (Figure 1A). Probably, the changes in NO emission under these conditions were below the detection limit of the NO analyzer.

**Figure 1:**
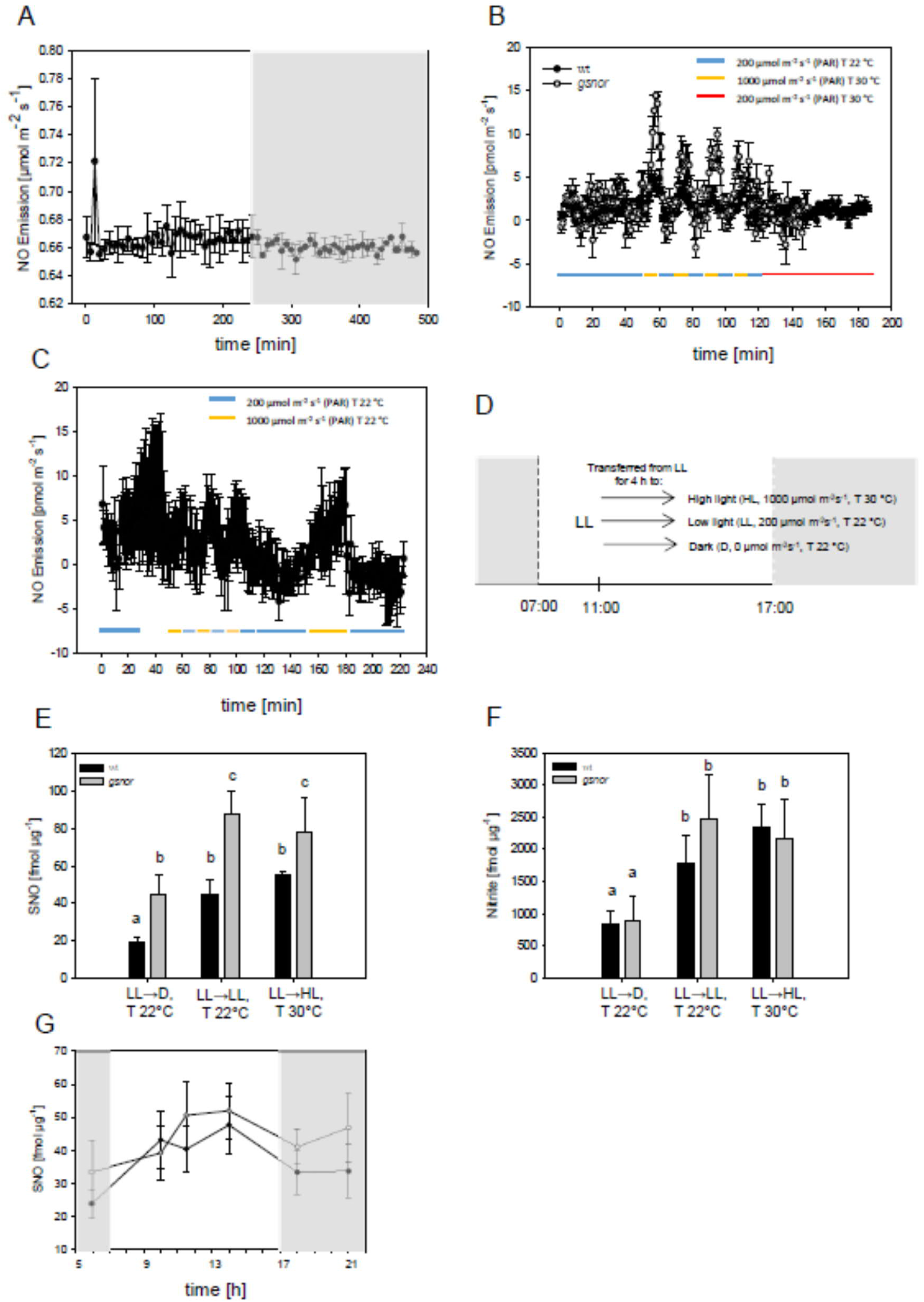
Light-dependent NO emissions and nitrite and S-nitrosothiol accumulation in Arabidopsis plants. (A-C) Single plants were put into an Arabidopsis cuvette and NO emission was measured by chemiluminescence using a ultra-high sensitive NO analyzer. Temperature and dark and light conditions were applied as indicated. PPFD, photosynthetic photon flux density (μmol photons m−2s−1). Shown are means ±SE of at least three independent experiments (N≥3). D) Four-week-old plants grown on soil at short day cycles (10/14 h light/dark, 20/17 °C) were transferred at noon (11:00) for 4 h to dark (D, T 22 °C), low light (LL, T 22 °C) or high light (HL, T 30 °C). Total S-nitrosothiol (E) and nitrite (F) levels were determined after 4 h. Shown is the mean +SE of three independent experiments (N=3). Letters are assigned to bars based on one-way ANOVA with Tukey’s post-hoc test. Two-way ANOVA results: in E) difference among light and temperature conditions – p=0.009, difference between wt and *gsnor* mutant – p=0.004. Pairwise comparisons were performed using the Holm-Sidak Test: D vs HL – p=0.020, D vs LL – p=0.014. In F) difference among light and temperature conditions – p=0.011. Pairwise comparisons: D vs HL – p=0.017, D vs LL – p=0.029. In G) total S-nitrosothiol levels of wt and *gsnor* plants were determined at 06:00, 10:00, 11:30, 14:00, 18:00 and 21:00 o’clock. The light period was from 07:00 to 17:00. Shown are the means ±SE of at least three independent experiments (N>=3).

In contrast, simulation of sunflecks, i. e. transient exposure to high light (HL, PPFD of 1000 µmol photons m^-2^s^-1^) in combination with increased temperature (T 30°C) led to a significant emission of NO in wt plants (approx. 5-fold) in comparison to low light/low temperature conditions (LL, 200 µmol photons m^-2^s^-1^, T 22°C) (Figure 1B). The increase in NO emission was higher in plants with loss of GSNOR activity (*gsnor*) (approx. 2-fold) in comparison to wt plants (Figure 1B). Increase of temperature alone did not enhance NO emission (Figure 1B), while high light intensity and constant temperature significantly increased NO emission (Figure 1C), demonstrating a link between light intensity and NO emission independent of temperature.

Since differences in NO emission could result from differences in stomata opening, endogenous SNO and nitrite contents were determined in Arabidopsis leaves. Plants were grown for four weeks under short day, low temperature (22°C) and LL conditions. After exposure for 4 h to LL, plants were transferred for 4 h to darkness (D, T 22°C) or high PPFD (HL, 1000 µmol photons m^-2^s^-1^, T 30°C) or kept under low PPFD (LL, 200 µmol photons m^-2^s^-1^, T 22°C) conditions (Figure 1D). Afterwards, SNO and nitrite contents were determined. In general, SNO content was higher in *gsnor* than in wt plants (Figure 1E). When kept under LL intensity, the SNO content in wt and *gsnor* plants was 45 and 88 fmol µg^-1^ protein, respectively. In both lines, the SNO level did not significantly increase when plants were transferred to high light intensity. However, plants transferred to darkness exhibited significantly lower SNO levels than plants kept under low or high PPFD intensities (wt: 45 to 19 fmol µg^-1^ protein, *gsnor*: 88 to 44 fmol µg^-1^ protein), further demonstrating a link between SNO formation and light intensity. The nitrite content is a frequently used option to display NO accumulation (He et al., 2004; Holzmeister et al., 2011). The nitrite levels under the different irradiation conditions correlated with the SNO contents for *gsnor* and wt plants (Figure 1F). Among the different PPFD levels, wt and *gsnor* plants did not significantly differ in their nitrite contents, but significantly lower nitrite levels were detected in wt and *gsnor* plants when transferred to the dark. Additionally, endogenous SNO levels of 4-weeks old plants grown under short day conditions (10/14 h light/dark, 20/17°C) were determined at different time points during the light and dark period (Figure 1G). A tendency to enhanced SNO concentration was observed under light, whereas lower SNO amounts were measured in the dark, further confirming a light-dependent accumulation of SNOs. Loss of GSNOR function resulted in slightly higher SNO levels in comparison to wt (Figure 1G).

### 2.2. GSNOR regulates histone acetylation in plants transferred to dark and different light conditions

Previously, we have demonstrated that exogenously applied NO donors and endogenously induced NO production results in enhanced histone acetylation (Mengel et al., 2017). Therefore, we analyzed whether light-dependent accumulation of NO/SNO also leads to chromatin remodeling in wt and *gsnor* plants. Four-week-old plants of both lines were exposed to different light conditions as mentioned above (see 2.1.), and global leaf levels of H3ac, H3K9ac and H3K9/14ac histone marks were quantified by immunoblotting using histone mark-specific antibodies (Figure 2A-D). Here, wt plants showed a tendency to continuously increase H3ac and H3K9/14ac levels from D to LL to HL conditions (Figure 2B and D). We observed significant approx. 2.5-fold and 2-fold increases in HL conditions compared to D conditions for H3ac and H3K9/14ac, respectively (Figure 2B and D). Moreover, the H3K9ac level tended to be higher in LL and HL conditions in comparison to D (Figure 2C, adjusted p-values 0.090 and 0.053, respectively). In *gsnor* plants, total H3ac, H3K9ac and H3K9/14ac levels were similar across the different light conditions and in most cases not significantly higher than the wt D level (Figure 2B-D). Thus, the correlation of histone acetylation with light intensity observed in wt was not present in *gsnor* plants. These data demonstrate that disturbed SNO homeostasis affects dark/light-dependent histone acetylation, suggesting a regulatory function of GSNOR (SNOs) in histone acetylation under these conditions.

**Figure 2:**
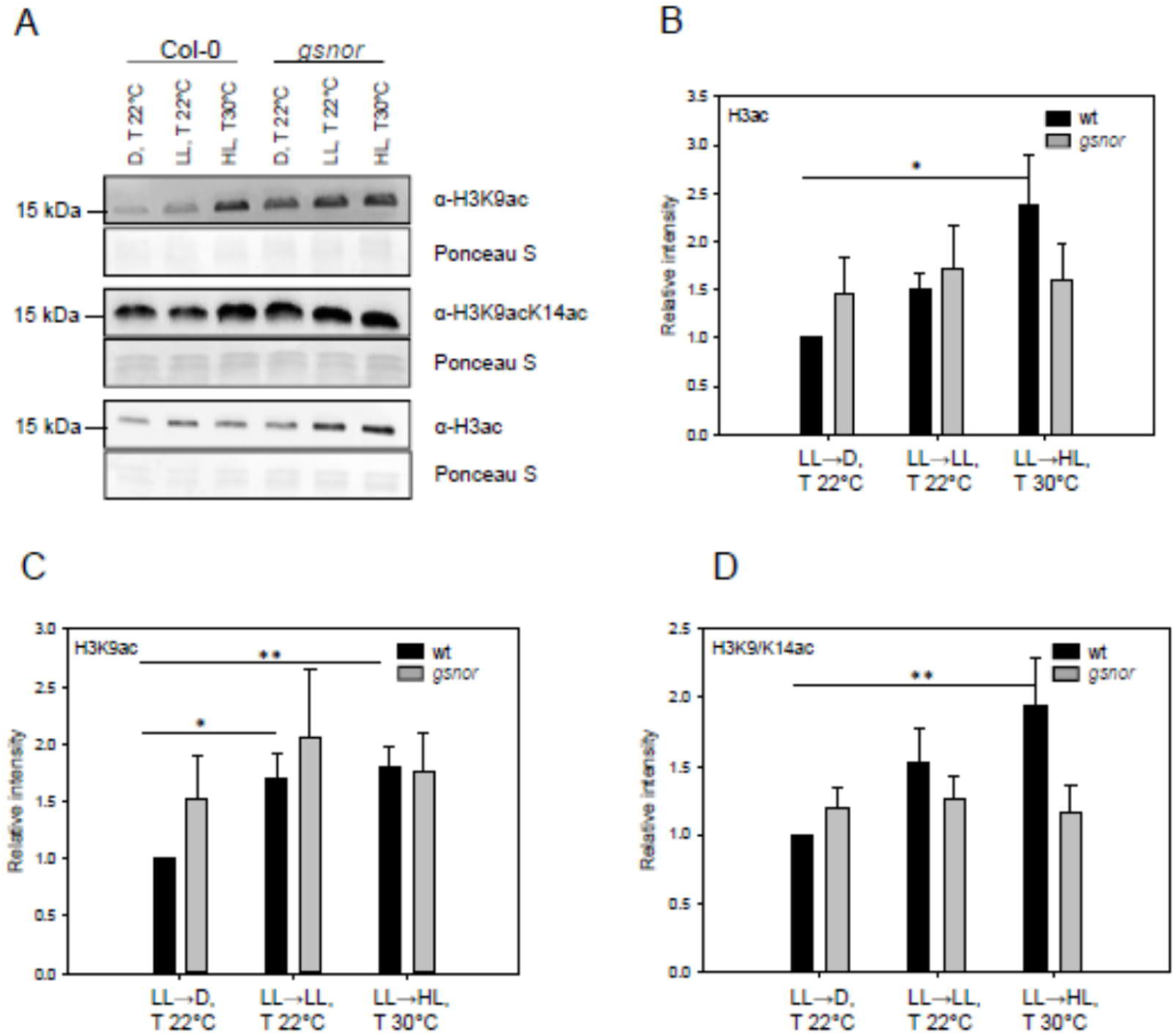
Different light conditions lead to altered H3 acetylation in wt and *gsnor* plants. Four-week-old plants grown on soil in short day (10/14 h light/dark, 20/17 °C) were transferred at noon (11 a.m.) for 4 h to dark (D, T 22 °C), low light (LL, T 22 °C) and high light (HL, T 30 °C). A) Histones were extracted, separated on a 12 % polyacrylamide gel and transferred onto a nitrocellulose membrane. The following antibodies were used for immunodetection of histone marks: acetylated-H3 (1:20000), acetylated-H3K9 (1:5000), acetylated-H3K9/14 (1:2000), acetylated-H4 (1:20000), and acetylated-H4K5 (1:10000). Anti-rabbit HRP (1:20000) was used as secondary antibody. B-D) Quantitative analysis of the immunodetected bands of the different histone marks. Signal intensity was determined with Image J software. Shown is the mean ±SE of at least five independent experiments (N≥5). Intensities are given relative to the histone acetylation level in wt under D conditions, which was set to 1. Significant deviations from this constant were determined by Holm adjustment after one-way ANOVA (**p≤0.01, *p≤0.05).

### 2.3. Identification of putative NO-sensitive HDAs

In Mengel et al. (2017), we already demonstrated that total HDA activity is reversibly inhibited *in vitro* by different NO donors (GSNO and SNAP), and *in vivo* by SA and INA, which both stimulate endogenous NO accumulation. However, it is still unclear which of the 18 different Arabidopsis HDAs are sensitive to NO. Members of the RPD3-like subfamily are the most promising candidates, since this subfamily includes HDA homologues to human HDA2. Mammalian HDA2 is S-nitrosated at Cys262 and Cys274 in response to NO resulting in chromatin remodeling in neurons and in dystrophic muscles (Colussi et al., 2008; Nott et al., 2008).

Comparison of the amino acid sequences of human HDA2 and Arabidopsis RPD3-like enzymes revealed that HDA6 is the closest homolog to human HDA2, with a sequence identity of 61 %. Both proteins contain seven conserved Cys, which are located within the HDA domain, including two Cys residues (Cys273 and Cys285 of AtHDA6) that correspond to the S-nitrosated Cys residues in mammalian HDA2 (Figure 3A). Interestingly, Cys273 of Arabidopsis HDA6 is a predicted target for S-nitrosation using the bioinformatic prediction tool GPS-SNO (Xue et al., 2010). Structural modeling of Arabidopsis HDA6 using the crystal structure of human HDA2 as template revealed strikingly similar 3D folds of both proteins (Figure 3B). In the structural model of Arabidopsis HDA6, Cys273 and Cys285 are located at the same positions as the S-nitrosated Cys262 and Cys274 in the 3D structure of human HDA2 (Figure 3B), indicating that both proteins exhibit a very similar microenvironment around the substrate binding site.

**Figure 3:**
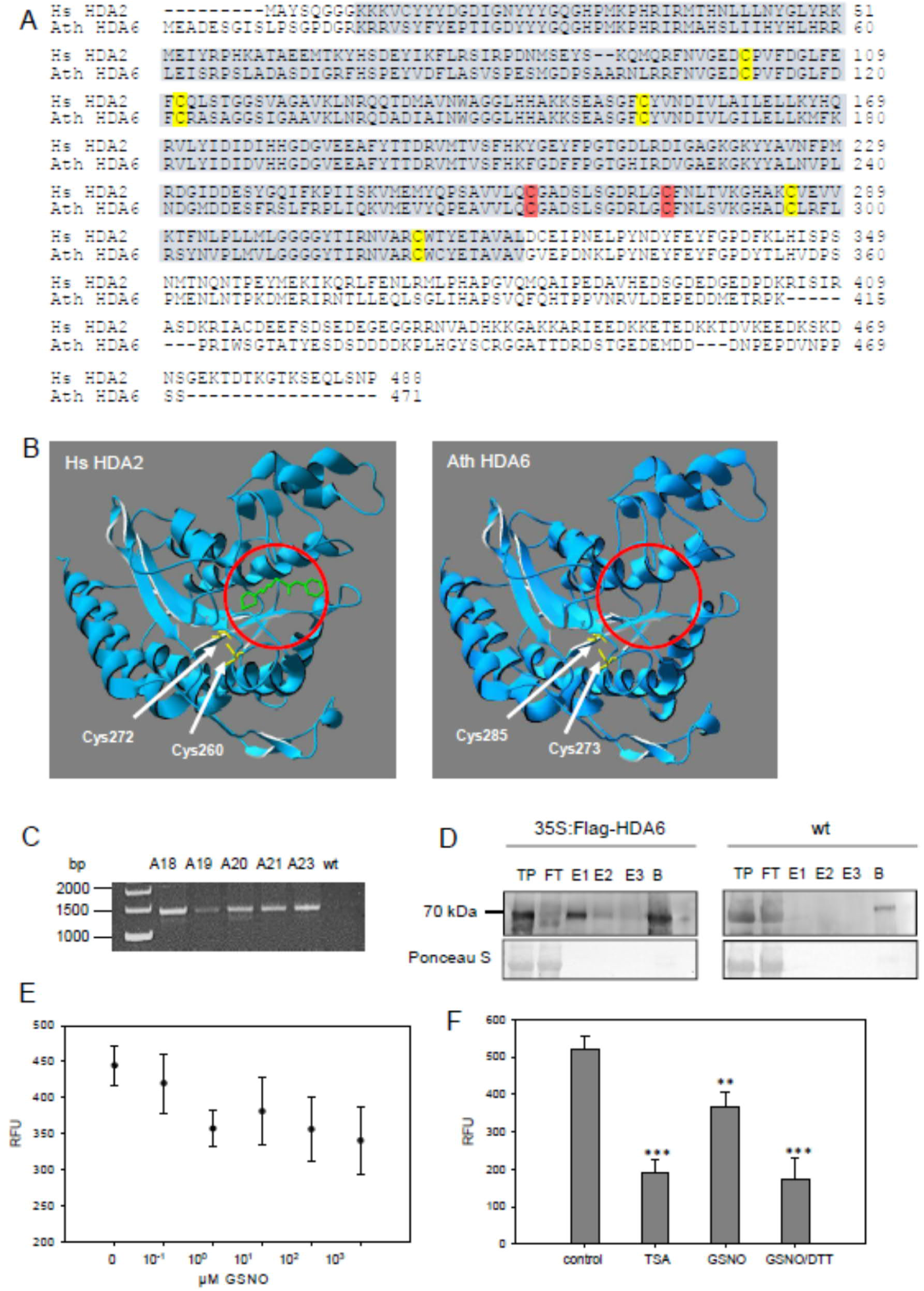
S-Nitrosation of Arabidopsis HDA6. A) Amino acid sequence alignment of human HDA2 and Arabidopsis HDA6 was performed using Clustal Omega. Cysteine residues that are S-nitrosated in human HDA2 are marked in red, other conserved cysteines are indicated in yellow. Histone deacetylase region is highlighted in blue. B) *A. thaliana* HDA6 displays a similar protein folding as human HDA2. The HDA domain of HDA6 (amino acids 18 – 386, Uniprot entry Q9FML2) was modelled using the SwissProt Modelling server with human HDA2 as a template (PDB code: 4LXZ). The 3D models were visualized with Swiss-PdbViewer. Cysteine residues that are S-nitrosylated in human HDA2 as well as the corresponding putative redox-sensitive cysteines of HDA6 are indicated. The bound HDAC inhibitor suberanilohydroxamic acid (green) in human HDA2 indicates its active center, which is highlighted in both enzymes with a red cycle. Recombinant FLAG-HDA6 was produced in *A. thaliana*. C) RT-PCR of transgenic 35S:FLAG-HDA6 Arabidopsis lines. Five 35S:FLAG-HDA6 containing lines A18-A21, and A23 were identified. cDNA of wt was used as a negative control. Predicted size of *FLAG-HDA6* is around 1470 bp. D) Immunoblot of in plants produced FLAG-HDA6. Total protein (TP) was extracted from 1g of the transgenic line A18 and wt and subjected to FLAG resin. Recombinant protein was eluted with 200 ng/ml Flag peptide for three times (E1-E3). TP, flow-through (FT) and E1-E3 were separated on a polyacrylamide gel and transfer onto nitrocellulose membrane. Anti-FLAG-tag antibody (1:1000) was used for immunodetection. Predicted size of FLAG-HDA6 is 57 kDa. Shown is one representative experiment of at least three replicates. E) Inhibition of FLAG-HDA6 activity by GSNO. The recombinant plant FLAG-HDA6 was incubated with 0.1 – 1000 μM GSNO for 20 min and its activity was determined. F) Activity of FLAG-HDA6 after treatment with 1 μM TSA, 1 mM GSNO and 1 mM GSNO/5 mM DTT. HDA activity was measured using Fluorogenic HDA Activity Assay. Shown is the mean ±SE of at least three independent experiments (N≥3). One-way ANOVA (DF=3; p<0,001) was performed with Holm-Sidak post-hoc test for each treatment group vs. the control group (FLAG-HDA6 activity), **p≤0.01, ***p≤0.001.

An *hda6* cell suspension line was generated to determine whether NO-dependent inhibition of total HDA activity is altered upon the knockout of HDA6. The *hda6 axe1-5* allele used to generate the cell culture contained an insertion resulting in a premature stop codon and the expression of a non-functional, C-terminally truncated version of the HDA6 protein (Murfett et al., 2001). Cell cultures exhibited similar growth kinetics and morphology to wt cells. Consistent with previous results (Mengel et al., 2017), wt cells showed a slight but significant increase in total H3ac level after GSNO treatment (500 µM), and a more pronounced, approximately 2.5-fold increase after TSA application (Supplementary Figure 1) (Mengel et al., 2017). In contrast, GSNO treatment of *hda6* cells did not result in an accumulation of acetylated H3 (Supplementary Figure 1). TSA treatment did not increase the rate of H3 acetylation either, indicating that HDA6 was the predominant TSA-sensitive HDA isoform in this cell culture system. Moreover, HDAC activity in wt nuclear extracts was sensitive towards NO, but TSA treatment could not completely abrogate HDAC activity (residual activity of 65 %; Supplementary Figure 2), indicating the presence of TSA insensitive HDACs (i.e. sirtuins). HDAC activity in *hda6* nuclear extracts was around 50 % lower compared to wt nuclear extracts (Supplementary Figure 2) and – consistent with the western blot results (Supplementary Figure 1) – was insensitive towards N-ethylmaleimide, a cysteine blocking compound (Supplementary Figure 3). These data make HDA6 a promising candidate to be a NO-sensitive HDA isoform.

To analyze if HDA6 can be S-nitrosated *in vitro* and if S-nitrosation indeed affects its activity, HDA6 protein was recombinantly produced in *E. coli* as His_6_-HDA6 and GST-HDA6 in BL21(DE3) cc4 (HDA6^E.coli^) - a strain which contains additional chaperones, helping to produce proteins with low solubility. However, no deacetylase activity could be measured for His_6_-HDA6 and GST-HDA6. We thus speculate that HDA6 might need certain posttranslational modifications or interaction partner(s) to function as an active histone deacetylase. We therefore produced recombinant FLAG-HDA6 in Arabidopsis. Presence of FLAG-HDA6 transcripts in transgenic lines were demonstrated by RT-PCR (Figure 3C). We purified recombinant FLAG-HDA6 protein and confirmed the presence of recombinant FLAG-HDA6 in transgenic lines with a predicted size around 55 kDa via immunoblot (Figure 3D).

Activity measurements demonstrated that recombinant FLAG-HDA6 was produced in a catalytically active form (Figure 3E). Treatment with increasing concentration of GSNO (up to 1000 µM) resulted in approx. 30 % inhibition of FLAG-HDA6 activity (Figure 3E), whereas 1 µM TSA reduced the activity by 65 % (Figure 3F). Surprisingly, the activity of GSNO-treated FLAG-HDA6 could not be restored by subsequent treatment with 5 mM DTT; in contrast, addition of DTT further inhibited HDA6 activity by 30 % (Figure 3F). Taken together, these data make HDA6 a promising candidate to be a NO-affected HDA isoform.

### 2.4. HDA6 regulates histone acetylation in plants transferred to dark and different light conditions

As demonstrated above, exposure to increasing light intensities enhanced NO emissions and SNO accumulation (Figure 1). Since HDA6 is inhibited by NO/SNO, we investigated whether biochemical function of HDA6 is required for regulating histone acetylation under dark and light conditions. Total H3ac, H3K9ac and H3K9/14ac levels in *hda6* knockout plants (*axe1-5*) were analyzed under the different light conditions described above (Figure 1D).

H3K9ac and H3K9/14ac are known substrates for HDA6 (Luo et al., 2017). As shown in Figure 4, total H3ac, H3K9ac and H3K9/14ac levels significantly increased from D to HL conditions in wt plants (Figure 4A-D). Interestingly, the acetylation levels in *hda6* (H3ac, H3K9ac and H3K9/14ac) – similarly to the acetylation levels in *gsnor* – did not follow this trend. H3K9/14ac levels showed a significant 3.5-fold increase in comparison to wt only in dark conditions. These data indicate that beside GSNOR activity, HDA6 activity is involved in modulating the chromatin structure especially in the dark and under low light intensities.

**Figure 4:**
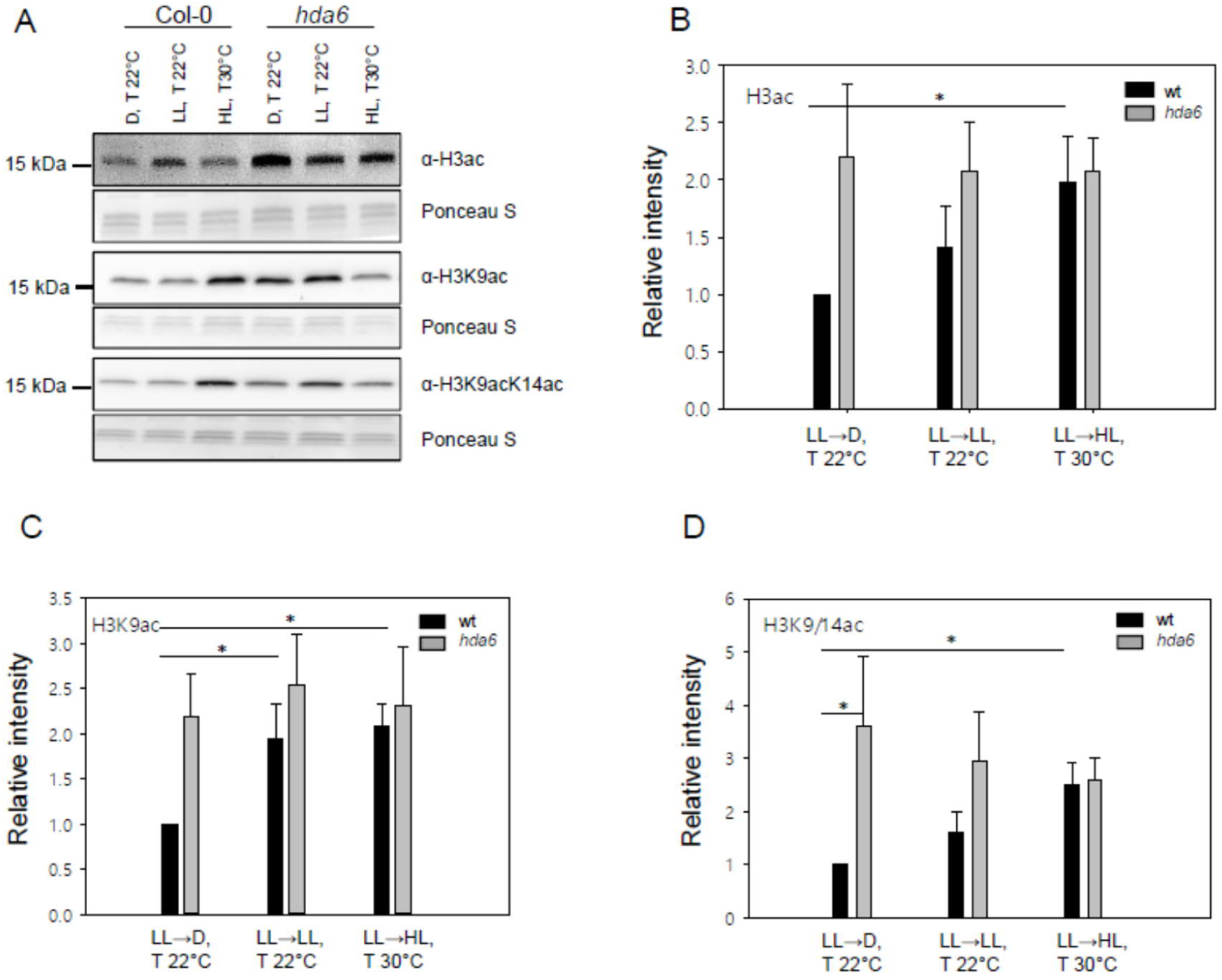
Different light conditions lead to altered H3 acetylation in wt and *hda6* plants. Four weeks old plants grown on soil at short day (10/14 h light/dark, 20/17 °C) were transferred at noon (11 a.m.) for 4 h to dark (D, T 22 °C), low light (LL, T 22 °C) and high light (HL, T 30 °C). A) Histones were extracted, separated on a 12 % polyacrylamide gel and transferred onto a nitrocellulose membrane. The following antibodies were used for immunodetection of histone marks: acetylated-H3 (1:20000), acetylated-H3K9 (1:5000), acetylated-H3K9/14 (1:2000), acetylated-H4 (1:20000), and acetylated-H4K5 (1:10000). Anti-rabbit HRP (1:20000) was used as secondary antibody. B-D) Quantitative analysis of the immunodetected bands of the different histone marks. Signal intensity was determined with Image J software. Shown is the mean ±SE of at least three independent experiments (N≥3). Intensities are given relative to the histone acetylation level in wt under D conditions, which was set to 1. Significant deviations from this constant were determined by Holm adjustment after one-way ANOVA (*p≤0.05).

### 2.5. Profiling of H3K9ac marks in light and dark conditions reveals differences of *gsnor* and *hda6* to wt

Under low light intensities and especially when plants were transferred to dark, histone deacetylation regarding H3, H3K9 and H3K9/14 depended on both GSNOR and HDA6 activity (Figure 2 and Figure 4). To identify chromatin regions regulated by GSNOR and HDA6 activity, we performed ChIP-seq using an anti-H3K9ac antibody. H3K9ac is a hallmark of active gene promoters (Karmodiya et al., 2012) and this histone mark shows a trend to higher abundance in *gsnor* (Figure 2C) and *hda6* (Figure 4C) mutants in comparison to wt under both dark and low light conditions. Four-week-old wt, *gsnor* and *hda6* plants were either exposed to low PPFD intensity or transferred to the dark for 4 h before a genome-wide light/dark-dependent H3K9ac profiling was performed by ChIP-seq. For all samples, the sequence reads aligned well with the *A. thaliana* genome, resulting in a total of 95.31-99.34 % aligned reads (Supplementary Table S1). After peak calling (Zhang et al., 2008), quantification and differential analysis were done to compare acetylation between light conditions and genotypes (Ross-Innes et al., 2012; Stark and Brown, 2019). Principle compound analysis (PCA) based on all the peaks demonstrated a good clustering of replicates (Figure 5A). Principle component 1 (PC1) shows light to dark effects for all genotypes (Figure 5A).

**Figure 5:**
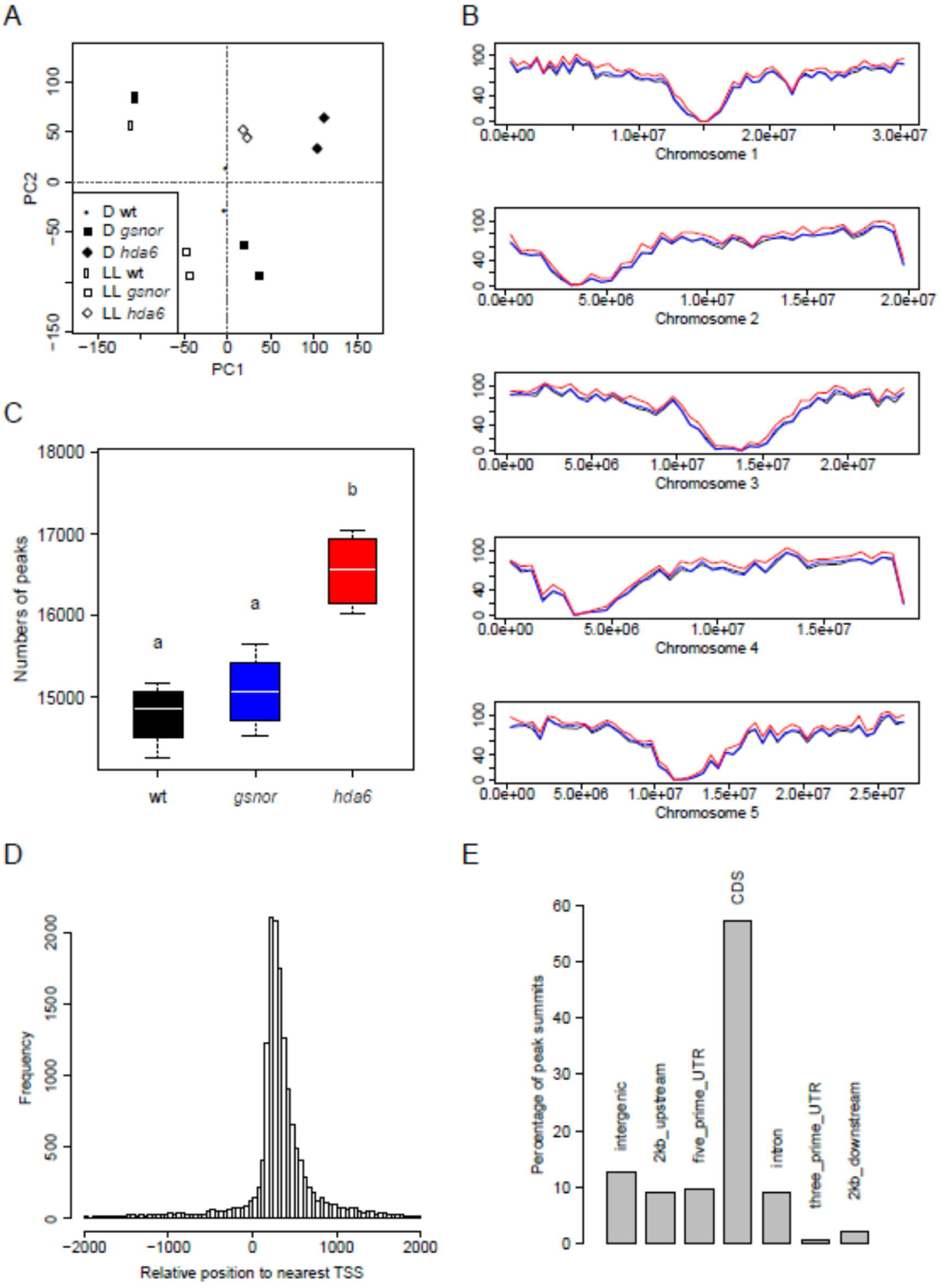
Characteristics of ChIP-seq samples. A) Principle component analysis (PCA). The projection onto the top two principal components (30% and 27% of variance, respectively) shows a clustering of biological replicates. Two independent ChIP-seq experiments were performed (N=2). B) Chromosomal location of H3K9ac peaks averaged for each line. Shown is the number of peaks in each 500 kb chromosomal bin of the *A. thaliana* genome. The centromeric and pericentromeric regions of each chromosome are characterized by very low number of peaks. Black: wt, blue: *gsnor*, red: *hda6*. C) Total number of identified peaks for wt, *gsnor* and *hda6*. Boxes show 25% and 75% quantiles, the white line represents the median and the whiskers indicate the extreme values. Lower-case letters mark groups that are statistically different (Kruskal Wallis test with posthoc Dunn test, p<0.05). D) Location of H3K9ac peaks relative to genes. Histogram of distances of peak summits to the closest annotated transcription start site (TSS). The distribution shows a maximum at 200 to 300 bp downstream of the TSS. E) Distribution of H3K9ac peaks according to the genomic region of the summit (relative to the closest TSS). CDS: coding sequence, UTR: untranslated region.

The highest density of H3K9ac peaks was found along the chromosome arms, whereas centromeric and pericentromeric regions were considerably less enriched in H3K9ac (Figure 5B). The number of H3K9ac peaks was significantly increased in the *hda6* mutant compared to wt and *gsnor* (Figure 5B and 5C). This hyperacetylation of DNA in *hda6* was observed throughout all chromosomes (Figure 5B). Most peaks were located 200 to 300 bp downstream of the closest TSS (Figure 5D). More than 93 % of all peaks were found within 2 kb upstream or downstream of a TSS.

Most of the peak summits (approx. 55%) are located within a coding sequence (CDS) and approx. 9 % were observed in five-prime untranslated regions (5’-UTR) and 2 kb upstream regions, respectively (Figure 5E). In total, we identified 16,276 H3K9ac peaks. Differences in H3K9ac between LL and D conditions were identified for each genotype (e. g., wt LL vs. wt D; adjusted p-value <0.05). All plant lines showed light-dependent acetylation changes with a positive effect preferentially on chloroplast and transport genes and a negative effect preferentially on stress response and transcription genes (Figure 6A, Supplementary Table S2, Supplementary Table S3). Peaks exclusively hyper- or hypoacetylated in wt or both mutants could also provide hints to the functions of HDA6 and GSNOR in the context of light stimulus-dependent histone acetylation. The wt was characterized by a hyperacetylation of stress responsive genes and a hypoacetylation of growth/development and chloroplast genes, whereas the mutants showed a hyperacetylation of genes involved in growth/development, carbohydrate metabolism and photosynthesis (Figure 6A, Supplementary Table S3).

**Figure 6:**
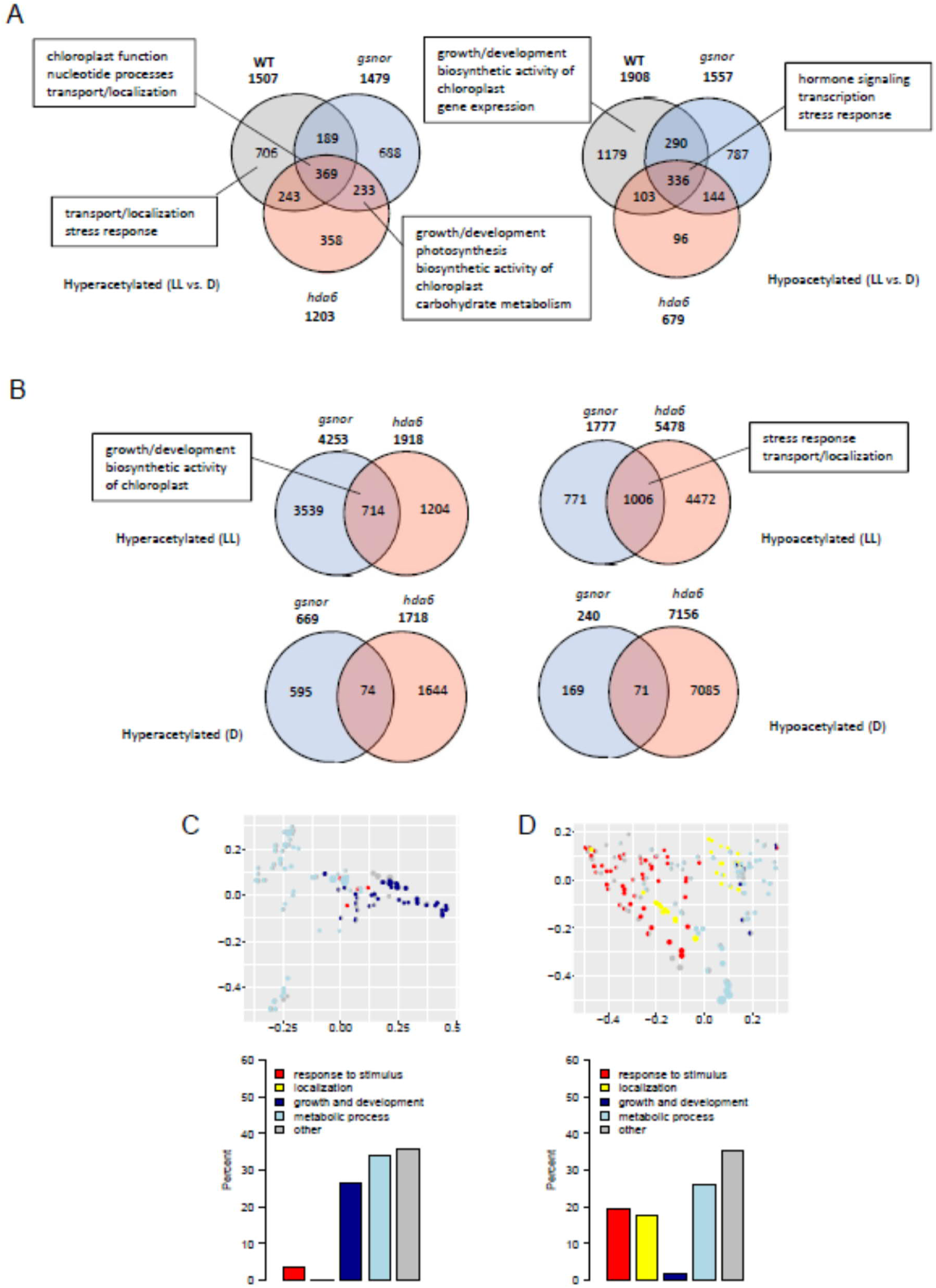
Differential acetylation in mutants. Venn diagrams of significantly changed peaks (adjusted p-value < 0.05) in LL vs. D (A) and mutant vs. wild type (B) comparisons. Boxes show major themes among significantly enriched GO terms (adjusted p-value < 0.05) for the respective partition. (C-D) Multi-dimensional scaling analysis of significantly enriched GO terms (adjusted p-value < 0.05) among the genes with closest TSS to significantly up-regulated (C) or down-regulated (D) acetylation peaks (adjusted p-value < 0.05) changed for both mutants vs. wild type under LL conditions. Only GO terms from the biological process ontology are shown in the plot. Each circle corresponds to an enriched GO term. Its size is proportional to the number of differentially acetylated genes (C: up, D: down) assigned to the GO term. The enriched GO terms are arranged in two dimensions such that their distance approximately reflects how distinct the corresponding sets of differential genes are from each other, i.e. neighboring circles share a large fraction of genes. Each enriched GO term is colored by its membership in the top level categories, which are grouped into five themes. If a GO term belongs to multiple top level terms, a pie chart within the circle indicates the relative fraction of each theme. The total distribution of themes across all enriched GO terms is depicted in the bar plot on the right.

### 2.6. GSNOR and HDA6 regulate H3K9ac of genes involved in growth/development, stress response, and localization in LL conditions

While the response to light already revealed differences between mutants and wt, a direct comparison of mutant and wt H3K9 acetylation under specific conditions will help to identify the basic functions of GSNOR and HDA6. H3K9ac peaks of *gsnor* and *hda6* were compared to H3K9ac peaks of wt plants both under LL and D conditions (e. g., *gsnor* LL vs. wt LL). The number of hyperacetylated H3K9ac peaks is higher in *gsnor* than *hda6* plants (Figure 6B). Remarkably, six times more H3K9 loci are hyperacetylated in LL in comparison to D conditions for *gsnor* (Figure 6B). In the *hda6* mutant, more H3K9 marks were hypoacetylated in comparison to wt than in *gsnor* (Figure 6B). Interestingly, both mutant lines share much more specifically hyperacetylated and hypoacetylated peaks in LL conditions in comparison to D conditions. 714 and 1,006 specifically hyperacetylated and hypoacetylated peaks, respectively, are shared under LL conditions with highly significant p-values for the overlap (2.7e-30 and 1.5 e-98, respectively), whereas 74 and 71 shared hyperacetylated and hypoacetylated peaks, respectively, were found in D conditions (Figure 6B).

To examine, which biological functions are shared by GSNOR- and HDA6-specific changes in chromatin acetylation, a GO term enrichment analysis was performed for the loci shared by both mutants. The corresponding genes of the hyperacetylated peaks (LL conditions) are enriched in GO terms, which mainly belong to growth/development (25%) and metabolic processes (>30%) including biosynthetic activity of chloroplast such as starch and pigment biosynthesis (Figure 6B-C, Supplementary Table S4). The genes identified within the hypoacetylated peaks (LL conditions) are enriched in GO terms related to stress response (approx. 20%), localization (approx. 20%) and metabolic processes (approx. 25%) (Figure 6B, D, Supplementary Table S4). In sum, these data suggest that GSNOR and HDA6 function is required to deacetylate particularly growth/development genes. Moreover, both enzyme functions promote acetylation of genes involved in stress response and localization.

### 2.7. Transcript profiling of wt, *gsnor* and *hda6* reveals gene regulation by light

Since H3K9ac is often found in actively transcribed promotors and coding sequences, we performed RNA-seq using the same experimental setup used for the ChIP-seq experiment. In all three genotypes around 6,000 genes are up-regulated or down-regulated (adjusted p-value <0.05) (Figure 7A, Supplementary Table S2). They share an overlap of 4,718 up-regulated and 4,598 down-regulated genes, which by design are independent of GSNOR and HDA6 function and are related to, e. g., chloroplast and ribosome functions (Figure 7A, Supplementary Table S5). In contrast, the 580 and 578 genes, which are exclusively up-regulated and down-regulated, respectively, in both mutants, depend on both enzyme functions. The up-regulated genes act in processes related to growth/development and transport/localization, whereas the down-regulated ones mainly functioning in stress response and transport/localization. Genes, that are exclusively up-regulated in wt, are enriched in GO terms also predominately related to stress response (Figure 7A, Supplementary Table S5). This is consistent with the increased acetylation of stress response genes in wt (Figure 6A). Taken together, our results suggest a GSNOR- and HDA6-dependent induction of stress responsive genes and repression of growth/development genes in response to light.

**Figure 7:**
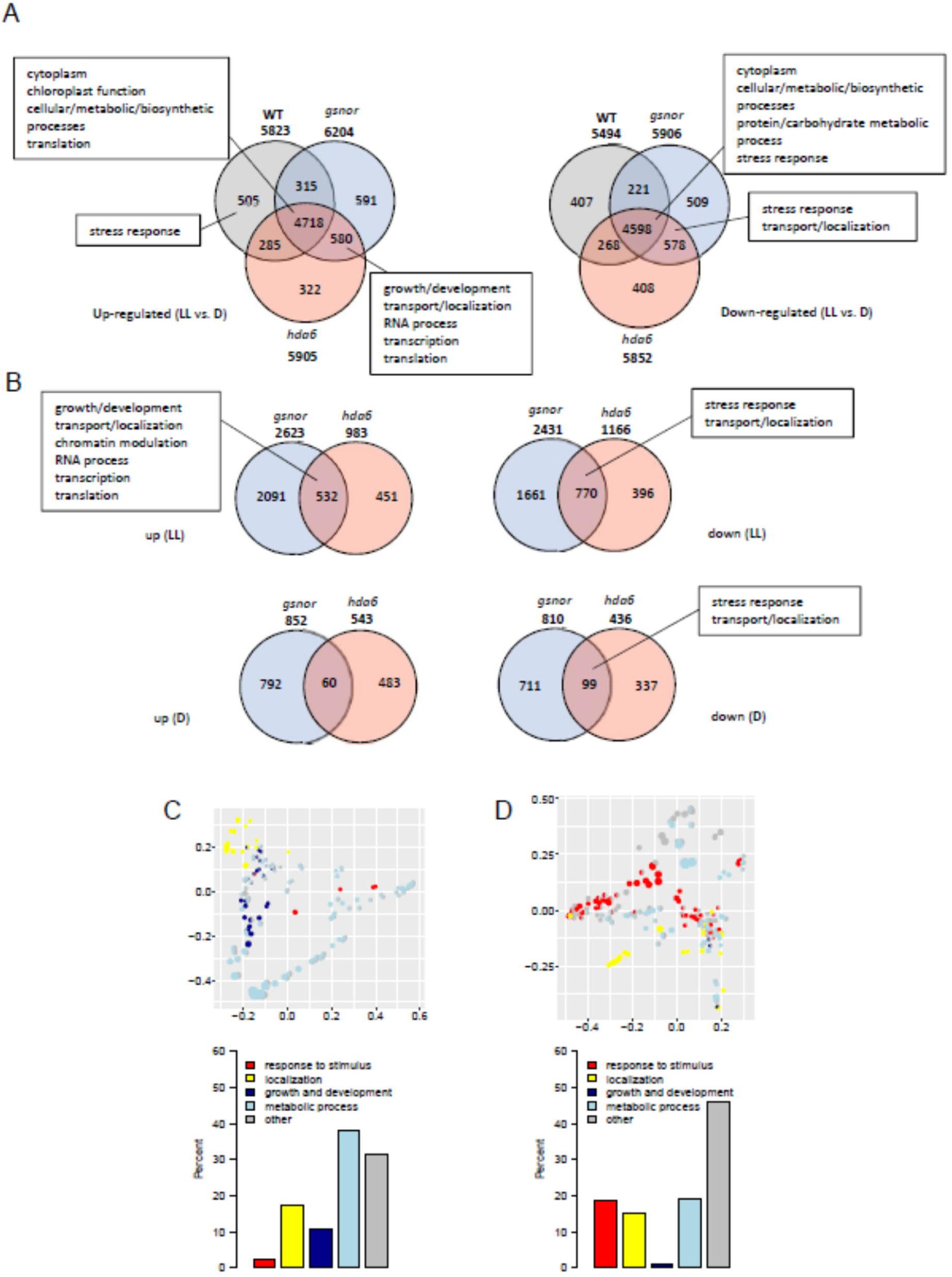
Differential gene regulation in mutants. Venn diagrams of significantly changed genes (adjusted p-value < 0.05) in LL vs. D (A) and mutant vs. wild type (B) comparisons. (C-D) Multi-dimensional scaling analysis of significantly enriched GO terms (adjusted p-value < 0.05) among the significantly up-regulated (C) or down-regulated (D) genes (adjusted p-value < 0.05) changed for both mutants vs. wild type under LL conditions. Only GO terms from the biological process ontology are shown in the plot. Each circle corresponds to an enriched GO term. Its size is proportional to the number of differentially regulated genes assigned (C: up, D: down) to the GO term. See Figure 6 for further details about the plots.

### 2.8. GSNOR and HDA6 regulate expression of genes involved in stress response, transport/localization and growth/development in LL conditions

To identify common regulatory functions of GSNOR and HDA6 activities in gene expression, the transcriptomes of *gsnor* and *hda6* were directly compared to wt, both under LL and D conditions. Similar to the acetylation data, in both genotypes more genes are differentially regulated under LL conditions than under D conditions (Figure 7B). Notably, the GO term enrichment results for the LL conditions share some overall trends with the ChIP-seq data (Figure 7C-D, Supplementary Table S6). While stress response functions are overrepresented among the genes down-regulated in both mutants, growth/development functions are only prominent among the up-regulated genes of both mutants. As a difference to the ChIP-seq results, many genes with transport/localization functions are up-regulated in both mutants.

### 2.9. Co-regulation between H3K9ac and gene expression in *gsnor* and *hda6* under LL conditions

To analyze the influence of H3K9ac on gene expression, ChIP-seq and RNA-seq datasets were integrated at the gene level for both mutants. Under LL conditions, the two mutants share 23 genes that show hyperacetylation and enhanced expression in comparison to wt plants. Interestingly, this group contains genes involved in growth/development, e. g. brassinosteroid biosynthesis (cytochrome P450 superfamily protein, AT3G50660), cell wall formation (glycosyl hydrolase family protein, AT1G78060), auxin biosynthesis (tryptophan aminotransferase related 2, AT4G24670), serine biosynthesis (D-3-phosphoglycerate dehydrogenase, AT4G34200) and histone modification (histone-lysine N-methyltransferase SETD1B-like protein, AT5G03670) (Figure 8A, Table, Supplementary Table S2). Under D conditions, only three genes are hyperacetylated and overexpressed in both mutants. One of these genes, AT5G16980 (Zinc-binding dehydrogenase family protein), has also shown up under LL conditions and is putatively involved in redox processes in plastids and cytosol.

**Table:**
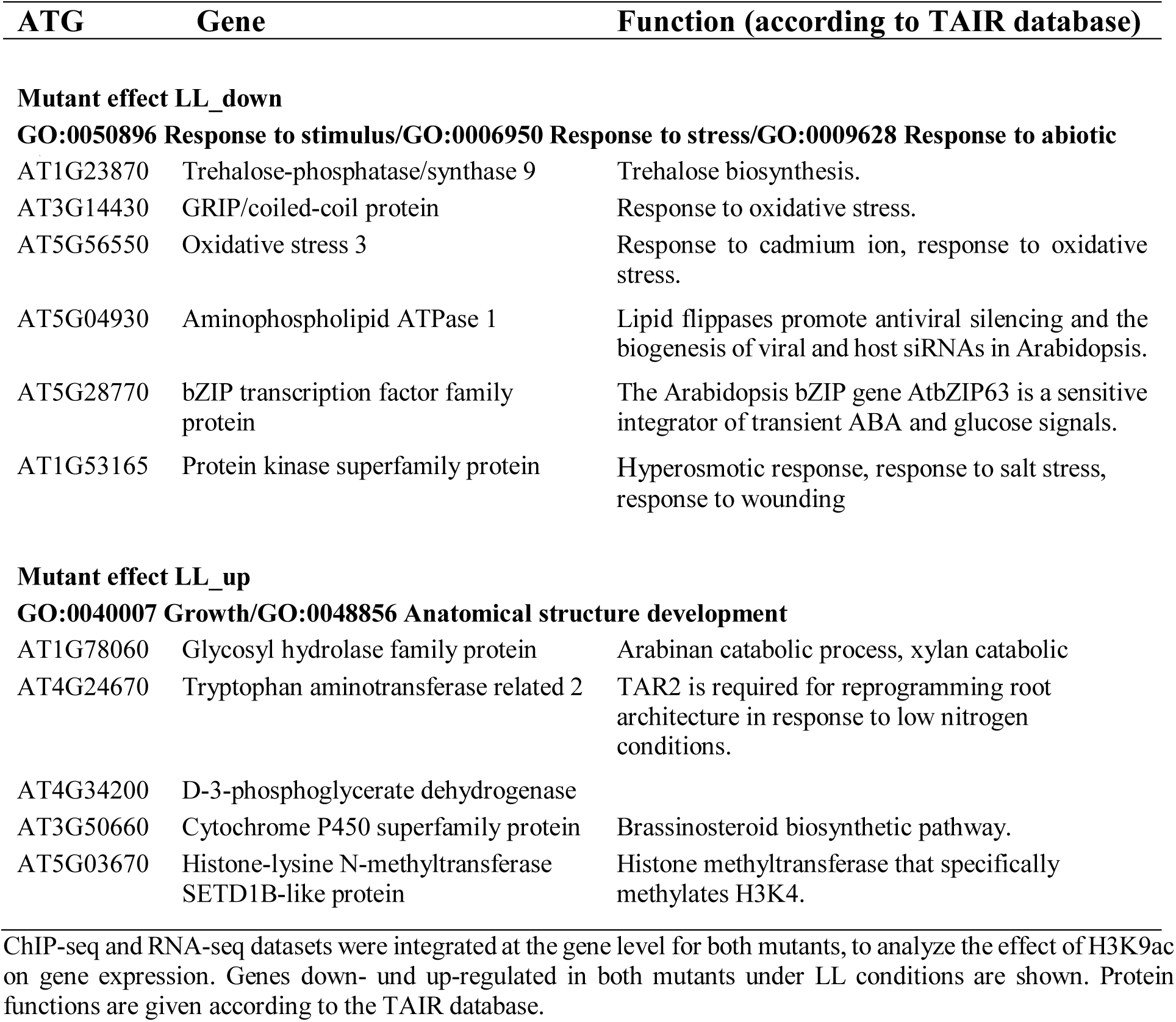
Selected genes showing correlated H3K9ac and gene expression.

**Figure 8:**
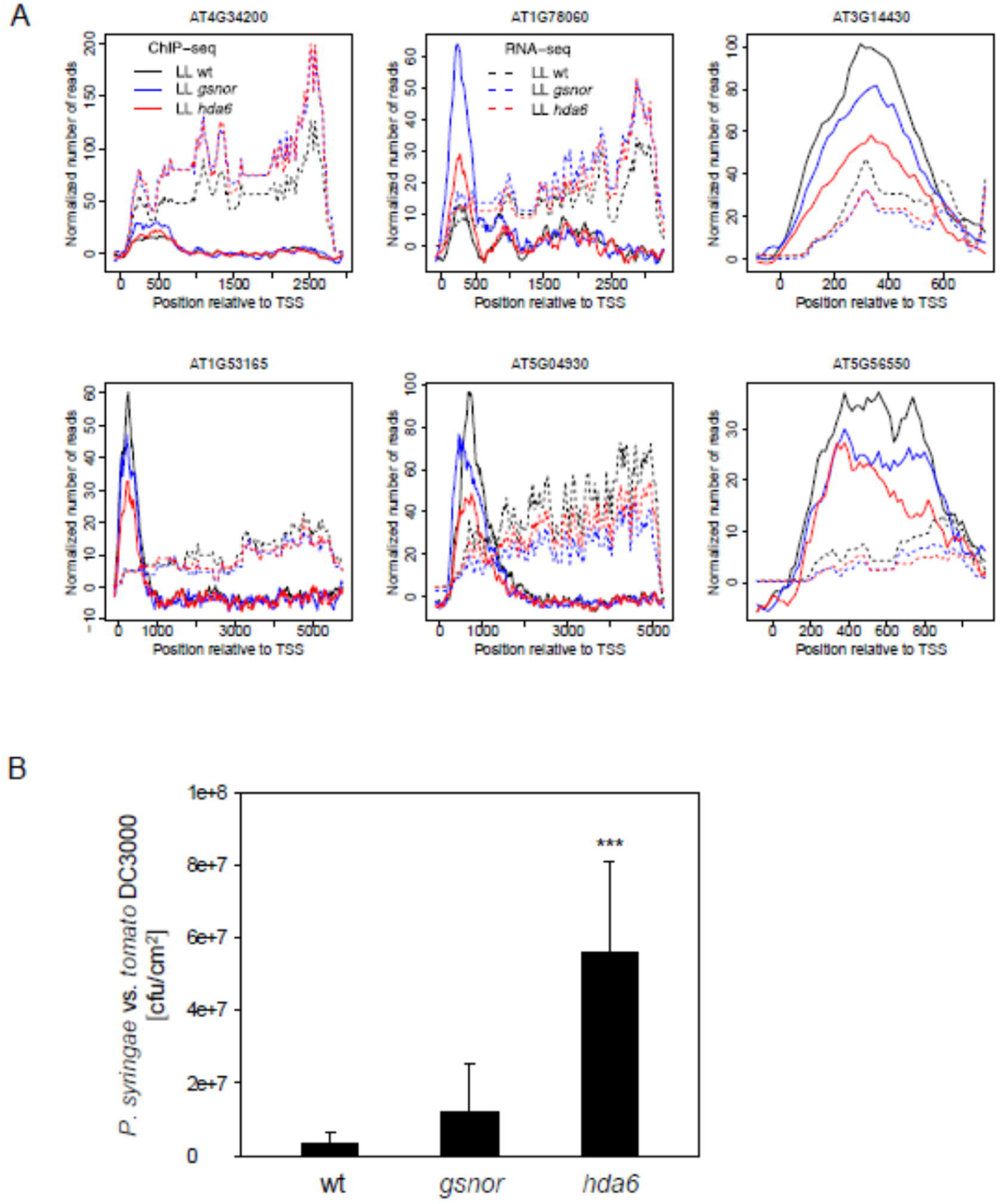
Comparative visualization of H3K9ac and gene expression. A) ChIP-seq and RNA-seq results of selected genes involved in growth/development (AT4G34200: D-3-phosphoglycerate dehydrogenase, AT1G78060: Glycosyl hydrolase family protein) and stress response (AT3G14430: GRIP/coiled-coil protein, AT1G53165: Protein kinase superfamily protein, AT5G04930: Aminophospholipid ATPase 1, AT5G56550: oxidative stress 3) are shown. B) Bacterial growth in infected wt and mutant plants. Shown is the mean ±SD for five biological replicates. Data were analyzed by one-way ANOVA with Dunnett’s posthoc test vs. wt (***, p <0.001).

In addition, 65 genes are hypoacetylated and less expressed in both mutants under LL conditions. These include several genes involved in abiotic and biotic stress response, e. g. RNA-binding KH domain-containing protein (AT1G14170), calcium-dependent protein kinase 2 (AT1G35670), protein kinase superfamily protein (AT1G53165), trehalose-phosphatase/synthase 9 (AT1G23870), aminophospholipid ATPase 1 (AT5G04930), oxidative stress 3 (AT5G56550), GRIP/coiled-coil protein (AT3G14430) and bZIP transcription factor family protein (AT5G28770) (Figure 8A, Supplementary Table S2).

The down-regulation of a significant amount of stress-related genes suggests a stress sensitivity of *gsnor* and *hda6* genotypes. To check, if the basal plant immunity system is affected, *gsnor* and *hda6* plants were infected with the virulent *Pseudomonas syringae* pv. *tomato* DC3000 (Figure 8B). Indeed, the virulent bacteria show a significantly stronger growth in *hda6* plants in comparison to wt plants, while *gsnor* plants displayed a non-significant increase (Figure 8B). The results demonstrate that HDA6 function is required for basal immune responses.

## 3. Discussion

### 3.1. Increasing light intensity enhances SNO accumulation and NO emission

NO is an important signaling molecule, which is involved in transcriptional regulation of many different physiological processes in plants, related to growth and development, abiotic and biotic stress response and photosynthesis (Huang et al., 2002; Kovacs et al., 2016; Kuruthukulangarakoola et al., 2017; Parani et al., 2004; Polverari et al., 2003). We observed an emission/accumulation of NO/SNO in the dark, which increased in light phases (Figure 1). GSNOR is responsible for controlling SNO homeostasis and loss of GSNOR function results in enhanced levels of SNO (Figure 1B, E-G). A similar observation was reported by other groups who demonstrated that endogenous NO production in Arabidopsis leaves exhibits a diurnal rhythm where the NO level was reduced by 30 % at night (He et al., 2004). Moreover, light-dependent NO release has been reported for tobacco leaves (Planchet et al., 2005). This indicates that light is an important trigger for intercellular accumulation of NO/SNO. NO production and emission under light could be based on light-triggered activation of nitrate reductase activity (Planchet et al., 2005; Riens and Heldt, 1992; Rockel et al., 2002). However, since the NO emission of nitrite reductase (NiR)-deficient tobacco leaves still increased in light, other factors besides these reductase activities might contribute to NO production, too (Planchet et al., 2005). For example, it could be possible that reduction of NO_2_^-^ to NO is related to photosynthetic electron flux where ferredoxin functions as electron donor. Reduction of NO_2_^-^ to NO is also possible in mitochondria of plants and animals in the presence of NADH (Kozlov et al., 1999; Stoimenova et al., 2007). However, this reaction is only observed under low oxygen conditions (Gupta et al., 2010). NO production in mammals correlated with the expression and activity of NOS which is triggered by light (Ko et al., 2013; Machado-Nils et al., 2013). Although NOS enzymes have not been found in higher plants yet, it was demonstrated that NO can be produced in chloroplast via a NADPH-dependent oxidation of L-arginine, which is the substrate of NOS. It has been shown that L-arginine is one the most common amino acids in chloroplasts and there available in nanomolar concentrations (Jasid et al., 2006). The synthesis of L-arginine is controlled by the photosynthetic light reaction, suggesting that oxidative NO production might also follow a circadian-like pattern. Stomata are usually open during the day (light) and closed at night (darkness), which might affect NO emission from leaves. However, light-dependent accumulation of endogenous SNOs and nitrite (Figure 1E-G) excluded that the observed light-dependent NO emission is just due to light-regulated stomata opening. In sum, the observed light-dependent NO/SNO accumulation/emission suggests a signaling function of this redox molecule in light-regulated processes. Moreover, since *gsnor* mutants showed increased NO emission and SNO accumulation under light (Figure 1B, E-G), *in vivo* GSNOR activity seems to have a regulatory function in light-dependent NO/SNO homeostasis.

### 3.2. Light-induced NO/SNO accumulation correlates with histone acetylation

Beside the direct modification of metabolic pathways and regulation of gene expression, NO can target the modulation of the chromatin structure, which is a less investigated regulatory mechanism. We observed a positive correlation between light intensity, NO/SNO accumulation and histone acetylation. The higher the light intensity, the higher the amount of accumulated SNO (and released NO) and the higher the levels of global H3ac, H3K9ac and H3K9/14ac in wt plants (Figure 1, Figure 2). Such a correlation was not observed in *gsnor* plants (Figure 2), suggesting a regulatory function of GSNOR activity (lower SNO level, denitrosation) in histone acetylation under these conditions.

There are several pieces of evidence indicating that SNO-induced histone acetylation is a result of the inhibition of HDA activity. First, 500 µM GSNO and SNAP reversibly reduce total HDAs activity by about 20 % in protoplasts and nuclear extracts (Mengel et al., 2017). Second, stimulation of endogenous NO production also inhibits the catalytic HDA activity in protoplasts (Mengel et al., 2017). Third, there are hints that the activity of at least some HDA isoforms are redox-regulated. Redox-sensitive Cys residues have been described in Arabidopsis HDA9 and HDA19. It is suggested that the oxidation of these two HDAs promotes their deactivation and therefore enhances histone acetylation and enables expression of associated genes (Liu et al., 2015). Redox regulation of HDAs has been already described in animals and humans. E. g. brain-derived neurotropic factor (BDNF) causes NO synthesis and S-nitrosation of human HDA2 at Cys262 and Cys274 in neurons. However, in this mammalian system S-nitrosation of HDA2 does not inhibit its deacetylase activity, but causes its release from a CREB-regulated gene promoter. Oxidation of HDA2 results in enhanced H3 and H4 acetylation at neurotrophin-dependent promoter regions and facilitates transcription of many genes (Nott et al., 2008). A different study reported about S-nitrosation of HDA2 in muscle cells of dystrophin-deficient MDX mice (Colussi et al., 2008). Although NO-sensitive Cys of this enzyme are not identified yet, it was shown that the enzymatic activity of muscle HDA2 is impaired upon NO donor treatment. Furthermore, recombinant mammals HDA6 and HDA8 have been reported to undergo S-nitrosation resulting in inhibition of their catalytic activity (Feng et al., 2011; Okuda et al., 2015). Moreover, HDA4 and HDA5, as parts of a large protein complex, migrate into the nucleus upon S-nitrosation of protein phosphatase 2A (Illi et al., 2008). Based on the studies mentioned above mammalian HDAs seems to play an important role in redox-signaling, (i) directly via NO or ROS production or (ii) indirectly by impairing HDA activities.

Similar mechanisms seem to be present in plants, too. Interestingly, Arabidopsis HDA6 share approx. 60 % amino acid sequence identity with mammal HDA2, which is redox-sensitive (Figure 3A). Both proteins contain seven Cys residues, which are located within the HDAs domain. NO/SNO-sensitive Cys residues of human HDA2 are conserved in Arabidopsis HDA6 and are located at similar position in the 3D structure of the proteins. The catalytic activity of purified *in-planta* produced FLAG-HDA6 is partially inhibited by GSNO (Figure 1E-F). Since S-nitrosation of HDA6 could be detected using the biotin switch assay, NO-mediated inhibition of HDA6 activity could be caused by PTM of cysteine residues. Surprisingly, the activity of GSNO-treated FLAG-HDA6 could not be restored by subsequent treatment with 5 mM DTT. In contrast, addition of DTT further inhibited HDA6 activity by 30%. Probably these quite strong reducing conditions resulted in loss of complex partners important for HDA6 activity or caused structural changes of the HDA6 protein.

H3ac, H3K9ac and H3K9/14ac levels tend to be higher in *hda6* plants in D conditions in comparison to wt plants (Figure 4B-D), while under HL conditions the acetylation levels of the *hda6* mutant and wt are similar. Interestingly, HDA6 controls light-induced chromatin compaction in Arabidopsis (Tessadori et al., 2009). The *hda6* mutant displayed a significant lower heterochromatin index under LL intensities (< 400 µmol m^-2^ s^-1^) than the corresponding wt plants. However, at HL intensities (> 500 µmol m^-2^ s^-1^) the heterochromatin index increased drastically in *hda6*, indicating a regulatory function of HDA6 in light-induced chromatin compaction. Since low and high histone acetylation levels correlate with compact and loose chromatin structure, respectively, our data confirm that HDA6 is involved in light-dependent chromatin modulation and make HDA6 a promising candidate to be a NO-affected HDA isoform. The proposed mechanism for the deacetylating function of GSNOR and HDA6 in D conditions is shown in Figure 9.

**Figure 9:**
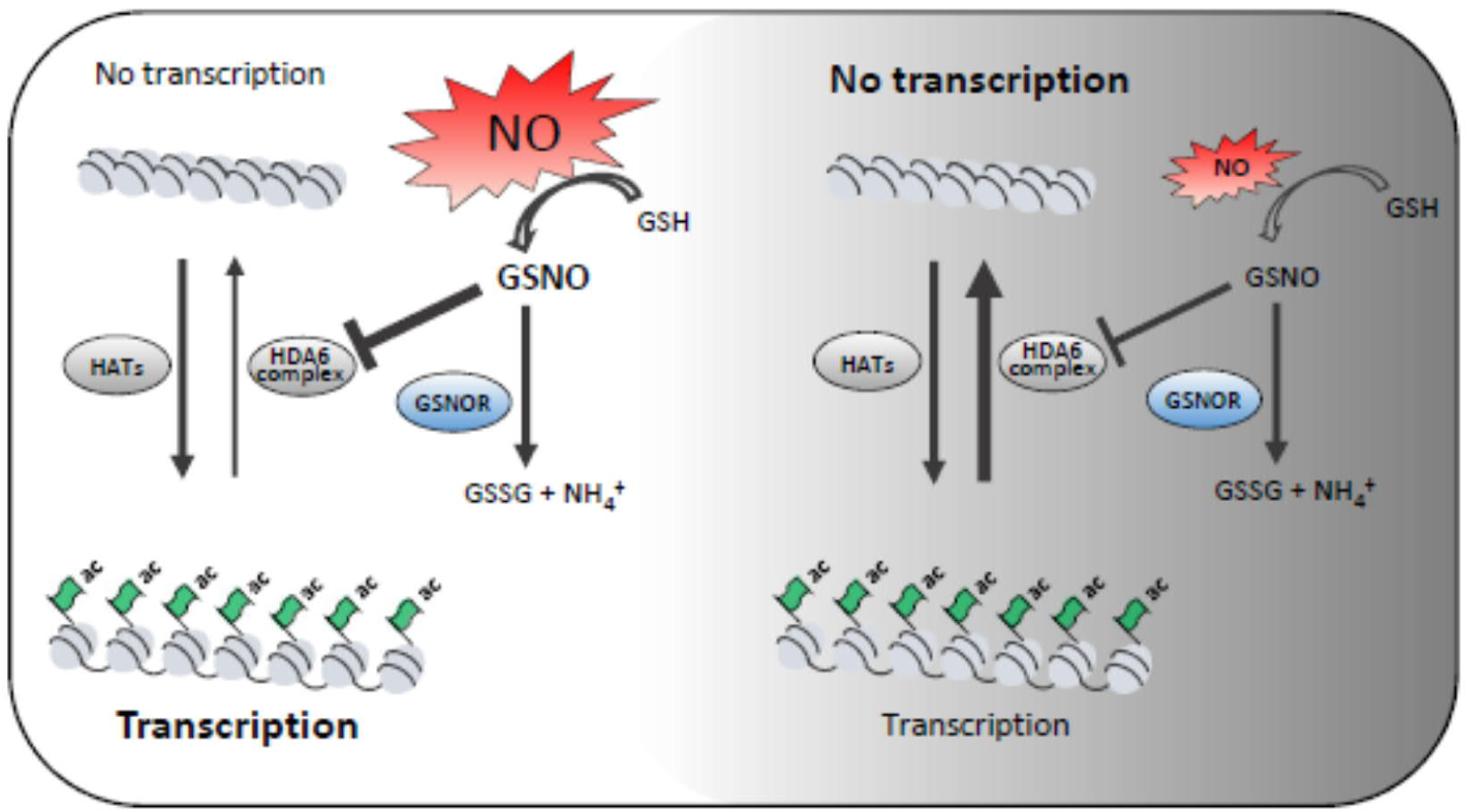
Schematic illustration of the regulatory function of NO on histone acetylation in light and dark conditions. Light-induced production of NO/GSNO results in enhanced inhibition of HDA6, increases histone acetylation, and gene transcription (left side). In dark condition, HDA6 activity is enhanced, because of less NO/GSNO production. As a consequence, histone acetylation and gene transcription are decreased. In both situations, GSNOR activity is required for fine-tuning the SNO levels.

### 3.3. Genome-wide profiling of H3K9ac in wt, *gsnor* and *hda6* shows enrichment close to TSSs

ChIP-seq analysis on H3K9ac was performed to examine the functions of GSNOR and HDA6 on chromatin structure in the dark and under light. 16,276 H3K9ac sites were found in the chromatin of Arabidopsis leaf tissue. Peaks mainly resided within gene-enriched areas and were almost depleted from centromeric and pericentromeric regions (Figure 5B). This observation is in line with reports from other plant species or other histone acetylation marks. For example, in the moss *Physcomitrella patens* H3K9ac and H3K27ac and in rice H3K9ac showed a strong enrichment in genic regions (He et al., 2010; Widiez et al., 2014). The genome-wide profiling of H3K9ac in Arabidopsis revealed that this histone modification is predominantly located within the regions surrounding the TSSs of genes and with a maximum at 200 to 300 bp downstream of the TSS (Figure 5D). This agrees with the distribution of H3K9ac in other plants as well as the distribution of other histone marks, e. g. analysis of different histone modification profiles in Arabidopsis revealed that most peaks are localized around 480 bp downstream of the TSSs, whereas peak position, shape, and length are independent of gene length (Ayyappan et al., 2019; Mahrez et al., 2016). Moreover, the preferential binding of transcription factors (∼ 86%) between − 1000 to + 200 bp from a TSS has been found in Arabidopsis (Yu et al., 2016). The *hda6* mutant displayed more acetylated regions than wt and *gsnor* throughout all chromosomes (Figure 5B-C).

### 3.4. GSNOR and HDA6 coordinate H3K9ac of genes involved in chloroplast function and growth/development

GO term analysis revealed that, under light conditions, *gsnor* and *hda6* share hyperacetylated genes related to chloroplast activity and growth/development (Figure 6A-C). These data suggest that GSNOR and HDA6 function is required to deacetylate these genes under light. Many plants produce and store metabolites and energy during the day, which are used for growth/development during night (Apelt et al., 2017; Graf et al., 2010). From this point of view, it makes sense that acetylation of genes involved in growth/development is reduced in light making these genes less accessible for the transcription machinery, whereas acetylation in the dark enables their transcription. Genes related to chloroplast function mainly concern starch, sulfur and terpenoid metabolism. Since the products of these genes are also required under light conditions, their reduced acetylation is surprising. However, since the acetylation levels of histones are a result of a fine-tuned interplay between acetyltransferases and histone deacetylases, GSNOR and HDA6 are probably just required to keep a balanced acetylation level of these genes. That is in line with the observation that expression of this set of genes is not changing significantly in both mutants in comparison wt.

The regulatory mechanisms of the deacetylating function of GSNOR under light are unknown. GSNOR activity lowers the level of GSNO and as consequence the level of S-nitrosated proteins. In this way, GSNOR is protecting HDA6 from SNO-dependent inhibition and keeping it active (Figure 9). However, our data do not rule out an additional effects of NO, e. g. activation of other HDAs or a reduced activity of distinct histone acetyltransferases. To get insight into the regulatory function of SNOs in chromatin modulation during light-dark switch, the S-nitrosylome under these conditions needs to be identified. In conclusion, according to the results obtained with the *gsnor* and *hda6* genotypes, both enzymes seem to play an important role in the light-dark (diurnal) regulation of histone acetylation.

### 3.5. GSNOR and HDA6 regulate H3K9 deacetylation and repression of genes involved in plant growth/development

Under light both mutants share several genes involved in growth/development, which show hyperacetylation and enhanced expression in comparison to wt plants, for instance histone-lysine N-methyltransferase SETD1B-like protein (AT5G03670). In Arabidopsis, 12 SET DOMAIN GROUP (SDG) containing histone methyltransferases are present, which are mainly involved in H3K4 and H3K36 methylation. These marks are active marks of transcription. So far, only a few genes of this gene family have been functionally characterized. SDG25, for instance, is involved in FLOWERING LOCUS C activation and repression of flowering (Berr et al., 2009). FLOWERING LOCUS C (FLC) is a key regulator of flowering, which negatively regulates downstream flowering activators such as FT and SOC1 (Helliwell et al., 2006). Consequently, high expression of FLC results in a late-flowering phenotype. Interestingly, in both *gsnor* as well as *hda6* mutants histone-lysine N-methyltransferase SETD1B-like protein acetylation and expression was increased and both mutants displayed a late-flowering phenotype (Kwon et al., 2012; Wu et al., 2008; Yu et al., 2011), assuming a flowering-activating role of GSNOR and HDA6. In *hda6* the late-flowering phenotype is likely due to up-regulation of FLC expression (Yu et al., 2011) (Supplementary Table S6). In contrast, for *gsnor* reduced or unchanged expression of FLC in comparison to wt plants is reported (Kwon et al., 2012); Supplementary Table S6), suggesting that GSNOR and HDA6 have different function in regulating flowering time.

Besides regulating the flowering time, GSNOR and HDA6 seem to have also important common regulatory functions in brassinosteroid biosynthesis. The gene encoding the cytochrome P450 superfamily protein (AT3G50660; DWARF4) was hyperacetylated and higher expressed in both mutants in comparison to wt plants. It encodes a 22α hydroxylase that is catalyzing a rate-limiting step in brassinosteroid biosynthesis (Choe et al., 2001). Brassinosteroids are phytohormones important for plant growth and development as well as for response to environmental stress. Mutants in the brassinosteroid pathway often display a dwarf phenotype (Kim et al., 2013; Li et al., 2001). Interestingly, *gsnor* displays a dwarf phenotype (Holzmeister et al., 2011; Kwon et al., 2012), although the key gene of brassinosteroid biosynthesis is upregulated, assuming that enhanced brassinosteroid biosynthesis is probably counteracting the dwarf phenotype resulted from GSNOR knockout. In sum, our results demonstrate that GSNOR and HDA6 are playing a role in negatively regulating histone acetylation and expression of genes involved in growth/development and chloroplast function.

### 3.6. GSNOR and HDA6 promote H3K9ac and expression of genes involved in plant stress response

Metabolic reprogramming in response to abiotic and biotic stress is governed by a complex network of genes, which are induced or repressed. A large set of stress-related genes is exclusively hyperacetylated in wt under LL vs. D conditions (Figure 6A). Light dependency of plant stress response has been investigated in the past in different contexts, e. g., circadian rhythm, day/night length and light composition (D’Amico-Damiao and Carvalho, 2018; Griebel and Zeier, 2008; Grundy et al., 2015; Sano et al., 2014). Various reports have shown that the plant signaling pathways involved in the responses to abiotic and biotic stresses are modulated by different types of photoreceptors controlling expression of a large fraction of abiotic stress-responsive genes as well as biosynthesis and signaling downstream of stress response hormones (Ballare, 2014; Jeong et al., 2010; Mazza and Ballare, 2015). For example, pathogen inoculations in the morning and midday resulted in higher accumulation of salicylic acid, faster expression of pathogenesis-related genes, and a more pronounced hypersensitive response than inoculations in the evening or at night (Griebel and Zeier, 2008). The observed plant defense capability upon day treatments seems to be attributable to the availability of a long light period during early plant-pathogen interaction rather than to the circadian rhythm. One might speculate, whether e.g. the light dependent flagellin 22-induced accumulation of salicylic acid (Sano et al., 2014) is related to H3K9ac. We observed light-dependent enrichment of the H3K9ac mark in many stress-related genes in wt in LL vs. D comparison (Figure 6A). Since H3K9ac is an activating histone mark, these genes might be prepared for expression and according to the RNA-seq data many stress-related genes displaying a higher expression in wt under light in comparison to darkness (Figure 7A).

Interestingly, GSNOR as well as HDA6 function seems to be involved in regulation of H3K9ac and expression of stress-related genes. Loss of GSNOR and HDA6 activity resulted in relative hypoacetylation and reduced expression of many stress-related genes (Figure 6B, D), suggesting that both enzymes are required to activate these stress-related genes. Given its HDA function, this means that those stress genes are specifically not targeted by HDA6. The loss of a distinct HDA function could result in activation of other HDAs or reduction of histone acetyltransferase activities, but this is not shown by our data (Figure 4). Rather, the increased overall number of acetylated regions might decrease the acetylation intensity at certain sites. Moreover, other still unknown factors could be involved in regulating histone acetylation. Indeed, indirect gene activating function has also been observed for other HDAs, e. g. for HDA5 (Luo et al., 2015), HDA9 (van der Woude et al., 2019) and HD2B (Latrasse et al., 2017).

Stress-responsive genes, which are hypoacetylated and down-regulated under light in both mutants include oxidative stress 3 (AT5G56550). Oxidative stress 3 is a chromatin-associated factor involved in heavy metal and oxidative stress tolerance (Blanvillain et al., 2009). It contains a domain corresponding to a putative N-acetyltransferase or thioltransferase catalytic site. Enhanced stress tolerance of *OXS3* overexpression lines and stress-sensitivity of *oxs3* mutant is favoring a role in stress tolerance. The nuclear localization of this protein supports a function as stress-related chromatin modifier protecting the DNA or altering transcription (Blanvillain et al., 2009).

Interestingly, acetylation and expression of trehalose-phosphatase/synthase 9 is also reduced in *gsnor* and *hda6* plants. Trehalose is a disaccharide composed of two glucose bound by an alpha-alpha (1 to 1) linkage and is often associated with stress-resistance in a wide range of organisms (Fernandez et al., 2010). Trehalose accumulation has been observed in plants under different stress situation, such as drought, heat, chilling, salinity and pathogen attack (Fernandez et al., 2010). Moreover, genes involved in detoxification and stress response are induced by exogenous application of trehalose (Bae et al., 2005a; Bae et al., 2005b; Govind et al., 2016) or by activating trehalose biosynthesis (Avonce et al., 2004).

The bZIP transcription factor family protein encodes for AtbZIP63, which is an important node of the glucose-ABA interaction network and may participates in the fine-tuning of ABA-mediated abiotic stress responses (Matiolli et al., 2011). The ABA signaling pathway is a key pathway that controls response to environmental stress.

The reduced acetylation and expression of stress-related genes in *gsnor* and *hda6* might be the reason for the susceptibility of both mutants against virulent *Pseudomonas syringae* vs. *tomato* DC3000 (Figure 8B). For *gsnor* plants, stress-sensitivity in context of pathogen infection, wounding, heat, cold, high salt, altered light conditions, and heavy metals has been described (summarized in (Jahnova et al., 2019)). Multiple roles in abiotic and biotic stress response are also known for HDA6 (Chen et al., 2010; Jung et al., 2013; Kim et al., 2017; Luo et al., 2012; Perrella et al., 2013; Popova et al., 2013; To et al., 2011; Wang et al., 2017). This underlines the importance of both proteins for effective stress response reactions. Previously, we published a putative link between NO/SNO and histone acetylation at stress-responsive genes (Mengel et al., 2017). We observed a SA-induced NO-dependent inhibition of total HDA activity and demonstrated a hyperacetylating function of exogenously applied GSNO at defense related genes. The temporally and spatially controlled production of NO as well as the presence or absence of NO-sensitive HDA-complexes could allow for the specific hyperacetylation of certain sets of stress-responsive genes (for instance S-nitrosation of a distinct HDA could specifically alter acetylation of salt-responsive genes). These NO-mediated histone acetylation changes could directly facilitate or enhance expression of the corresponding stress-related genes.

Emission of NO and SNO level was higher under light compared to darkness (Figure 1B-C, 1E-G). Although GSNOR inhibits the activity of recombinant HDA6 (Figure 3E-F), their effects on stress-related genes are probably indirect. In addition, other HDAs and histone acetyltransferases can be involved in regulation of histone acetylation at stress-responsive genes. It was demonstrated that HDA19 plays an essential role in suppressing SA-biosynthetic genes and PR-genes during unchallenged conditions by deacetylation of the corresponding promoters (Choi et al., 2012). After pathogen attack, histone acetylation at these regions increased suggesting a reduction of HDA19 activity or alternatively an activation/recruitment of histone acetyltransferase activity. Furthermore, in Arabidopsis the plant-specific HD2B is binding to genes involved in defense response in untreated plants, whereas after flg22 treatment mainly genes involved in plastid organization are targeted by HD2B (Latrasse et al., 2017).

All these observations highlight the importance of a fine-tuned switch between growth and development on one side and stress response on the other side. In this context, GSNOR and HDA6 seem to play a key role in coordinating histone acetylation and expression of stress-related genes and genes involved in growth/development to reduce plant growth/development and to allow a successful stress response (Figure 10). On the other side, the coordinating function of these enzymes and NO could be a promising target to modify plant metabolism to mitigating the negative effects of stressful environment on plant performance and productivity. Moreover, our study shows that, in addition to the known suppressive effects of HDAs, HDA6 has also indirect positive effects on transcription and interestingly, GSNOR activity seems to be involved in this process of switching the metabolism from growth and development to stress response. In sum, it appears that NO coordinates histone acetylation and expression of genes involved in growth/development and stress response.

**Figure 10:**
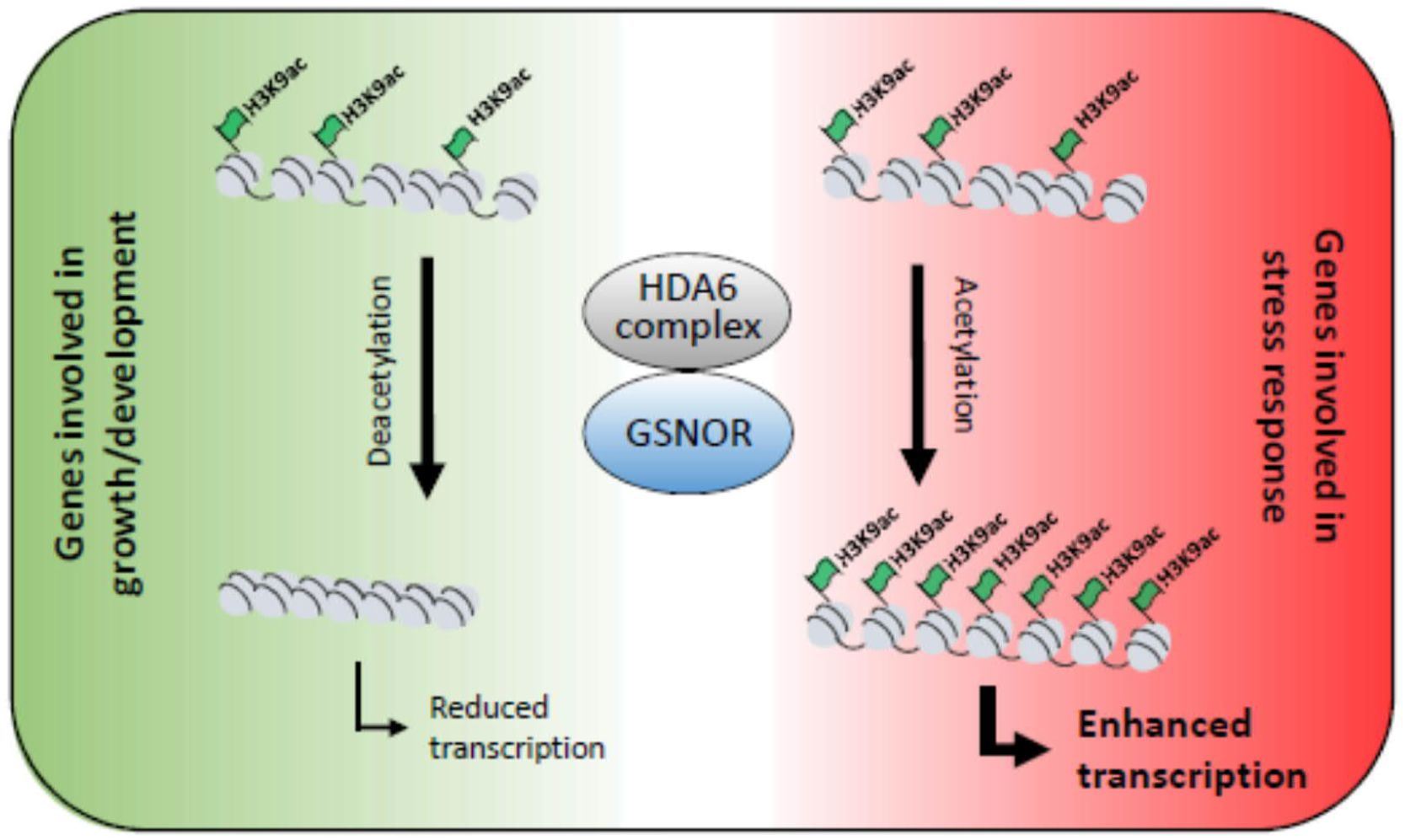
GSNOR and HDA6 differentially modulate H3K9ac of genes involved in growth/development and stress response. GSNOR and HDA6 act in similar pathways responsible for the regulation of an identical set of growth/development related genes as well as stress-related genes. While GSNOR and HDA6 function is required for deacetylation and repression of genes involved in growth/development, both enzymes are also involved in acetylation and enhanced expression of stress responsive genes, suggesting that GSNOR and HDA6 function as molecular switch between both physiological processes.

## Acknowledgments

We thank Elke Mattes, Lucia Gößl and Rosina Ludwig for excellent technical assistance. This work was supported by the Bundesministerium für Bildung und Forschung (BMBF).

## Author Contributions

Conceptualization, A.A. and C.L.; Methodology, A.A., B.W., A.G., and A.Al.; Investigation, A.A., P.H., and A.M.; Formal Analysis, E.G, A.A. and P.H.; Writing – Original Draft, A.A. and C.L; Writing – Review & Editing, A.A., C.L., E.G., B.W., A.Al., C.B., J.-P.S., and J.D.; Supervision, C.L., C.B., and J.-P.S.

## Materials and Methods

### Plant lines, cultivation

*A. thaliana* wild type Col-0, *gsnor1-3* [GABI-Kat 315D11; *gsnor*; *GSNOR-KO*], axe1-5 (*hda6*; *HDA6-ko*) were cultivated on soil mixed in ratio 1:5 with sand. The *hda6 axe1-5* allele used to generate the cell culture contains an insertion resulting in a premature stop codon and the expression of a non-functional, C-terminally truncated version of HDA6 (Murfett et al., 2001). Plants were grown under short day (10 h light/14 h dark and 20 °C/16 °C, respectively). The relative humidity during the day and night was 50 %. Light intensity in both conditions was approx. 100 to 130 µmol photons m^-2^ s^-1^ PPFD.

### Extraction of nuclear proteins

Nuclear proteins were extracted from Arabidopsis cell culture or seedlings according to the protocol of (Xu and Copeland, 2012) with small modifications. Approx. 0.5 - 0.6 g of grinded Arabidopsis tissue or cell culture were homogenized in 3 ml of LB buffer and filtered through two layers of miracloth and 40 µm nylon mesh sequentially. The homogenate was centrifuged for 10 min at

1500 g and 4 °C. The supernatant was discarded and the pellet was resuspended in 3 ml of NRBT buffer and centrifuged as described above. This step was repeated two more times or until the green color is gone (chloroplast contaminations). The triton X - 100 was removed from the nuclei pellet by washing it in 3 ml of NRB buffer. If the nuclei were not used immediately, they were resuspended in 400 µl of NSB buffer, frozen at liquid nitrogen and stored at - 80 °C.

Two methods were used to break a nuclear envelope and solubilize proteins. For quick detection of nuclear proteins by western blot, nuclear pellet was resuspended in 50 µl of Laemmli buffer, heated for approx. 10 min at 95 °C and centrifuged for 15 min at maximal speed. Protein concentration was measured using a RC DC protein assay (Biorad, Cat No 5000121). The second method was based on the sonication procedure using micro tip MS 72 (Bandelin, Cat No 492). The nuclei pellet was resuspended in approx. 300 µl of NPLB buffer and sonicated for 30 sec, step 3 and 20 – 40 %. The sonication step was repeated in total 5 times with approx. 1 min break in between. Protein concentration was measured using a Bradford reagent (Biorad, Cat No 5000006).

### Preparation of histones

Histone proteins were extracted either from in liquid grown seedlings or from leaf tissue with a Histone Purification Kit (Active Motif, Cat No. 40025) using manufacturing instruction with some modifications. 0.5 – 0.6 g start material were ground to a powder and incubated for 2 h with 2.5 ml extraction buffer on a rotating platform at 4 °C. The extracts were centrifuged at 4 °C for 10 min at maximal RCF. Afterwards the supernatants were transferred to PD 10 columns (GE Healthcare, Cat No. 17085101), which prior were equilibrated two times with 3.5 ml pre-cooled extraction buffer. The proteins were eluted with 3.5 ml extraction buffer. The eluates were neutralized with 1/4 volumes of 5 x neutralization buffer (0.875 ml) to reach a pH of 8. Purification of core histones was the same as in the instruction following the buffer exchange procedure using Zeba spin desalting columns 7K MWCO (Thermo Fisher, Cat No. 89882). Columns were prepared by adding three times 300 µl dH2O with a Protease Inhibitor EDTA-free tablet (Roche, Cat No. 04693132001). 100 µl of purified core histones were added to the column and centrifuged for 2 min at 1500 RCF. Histone amount was measured by NanoDrop 1000 at 230 nm.

### SDS - PAGE

Protein extracts were equally loaded on a precast 12% polyacrylamide (Biorad, Cat No 4561044) or self-made gel and subjected to a sodium dodecyl sulfate-polyacrylamide gel (SDS-PAGE) using a Mini-PROTEAN® Electrophoresis cell (Biorad, Cat No 1658002EDU). Gels were run at 130 V for approx. 60 min in 1 x running buffer. After separation of proteins a gel was either stained for 30 min with Coomassie brilliant blue solution or further used for western blot.

### Western blot

Proteins were transferred to a nitrocellulose membrane (Abcam) using a semi - dry western blot system. Pre-wet membrane and gel were sandwiched between whatman papers that were pre-soaked before in a transfer buffer. A transfer was performed for 45 min at room temperature. A flow rate of electric charged was dependent on length (L), width (W) and amount (n) of membranes and was calculated as follows: mA=L x W x 2.5 x n. An efficient transfer of proteins was determined by staining a membrane with Ponceau S solution (Sigma - Aldrich, Cat No 6226-79-5). Afterwards a membrane was incubated for 1 h in a blocking buffer shaking at room temperature followed by binding with primary antibody in 5 % BSA/TBS-T buffer overnight at 4 °C. A membrane was washed three times for 5 min with 1 x TBS-T buffer and incubated for 1 h at room temperature with horseradish peroxidase (HRP) - linked secondary antibody in 5% BSA/TBS-T buffer. A membrane was washed first once with 1 x TBS-T and two times with 1 x TBS buffer. The signal was developed using Western lightning plus-ECL chemiluminescence substrate (PerkinElmer, Cat No NEL105001EA).

### Recombinant expression and purification

The vector carrying a N-terminally FLAG-targeted HDA6 (pEarlyGate202/HDA6) was transferred to DH5α followed by electroporation of GV3101 pMP90. Transgenic Arabidopsis lines overproducing 35S:FLAG-HDA6 were generated by floral dip method as described above. Homozygous lines were selected and used for further studies. Plants expressing recombinant FLAG-HDA6 were harvested three weeks after sowing. For analytical studies around 4 g of ground material were used. Protein extracts were prepared in two volumes (approx.8 ml) of CelLyticP buffer (Sigma-Aldrich, Cat No C2360) with 1 % of a Protease Inhibitor EDTA-free tablet (Roche, Cat No. 04693132001) by rotating for 1 h at 4 °C. Extracts were filtrated trough miracloth (Millipore, Cat No 475855-1R) followed by 15 min centrifugation at 6000 x g and 4 °C. 60 µl of Flag-targeted beads (Sigma-Aldrich, Cat No A2220) were equilibrated with TBS buffer according to the manufacturer’s instruction and added to the extracted proteins. A binding of recombinant protein to the beads were performed at 4 °C rotating for 4 h. Afterwards the resin was centrifuged for 30 sec at 8200 x g and supernatant was discarded. The beads were washed tree times with TBS solution and FLAG-HDA6 was eluted with 200 ng/µl of Flag-Peptide (Sigma-Aldrich, Cat No F3290) by incubation the resin with synthetic peptide rotating for 30 min at 4 °C.

### Measurement of HDA activity

HDA activity was measured using a commercially available EpigenaseTM HDAC Activity/Inhibition Direct Assay Kit (Epigentek, Cat No. P-4035-48) according to the manufacturer’s instruction. 3-17 µl of purified Flag-HDA6 per well were treated with chemicals such as GSNO, GSH, TSA, DTT and incubated with 50 ng of substrate for 90 min at RT. HDA-deacetylated product was immuno-recognized and the fluorescence at 530Ex/590Em nm was measured in a fluorescent microplate reader (Tecan infinite 1000). The RFU values were directly used for relative quantification of HDA activity. HDA activity was also measured according to (Wegener et al., 2003). 3-17 µl of purified Flag-HDA6 per well were first treated with GSNO or TSA for 30 min in the dark at RT followed by incubation with DTT (if it was required) for another 30 min. The HDA reaction was started by adding 200 µM of HDA-substrate (Boc-Lys(Ac)-MCA) in 25 µl of HDA buffer followed by 60 min incubation at 37°C. The reaction was stopped by adding 45 µl of 2 x Stopping solution containing 10 mg/ml trypsin and 1 µM TSA. The mixture was incubated for an additional 20 min at 30°C to ensure the tryptic digestion. The release of 7-amino-4-methylcoumarin (AMC) was measured by monitoring of florescence at 380Ex/460Em nm.

### Nitrosothiol and nitrite measurement

S-Nitrosothiols and nitrite were measured using Sievers Nitric Oxide Analyzer NOA 280i (GE Analytical Instruments). The method is based on reduction of SNOs and nitrite to NO, that is further oxidized by ozone to NO_2_ (excited state) and O_2_. On the way to the ground state NO_2_ emits chemiluminescence which can be measured by photomultiplier. Approx. 300 – 500 mg of plant tissue were homogenized in the same volume of PBS solution and incubated for 20 min rotating at 4 °C. Protein extracts were separated from plant debris by centrifugation for 15 min at maximal speed. 20 – 100 µl of analyte were injected into triiodide solution. For the detection of SNO content sulfanilamide (1:9) was additionally added to protein extracts to scavenge nitrite and 200 µl were injected. Every measurement was performed in duplicates. A standard curve was created with sodium nitrite.

### Measurement of NO emission

NO emission was measured from 3.5 – 4-week old Arabidopsis plants using a CLD Supreme chemiluminescence analyzer (ECO PHYSICS). The purified measuring gas with a constant flow of 600 ml/min was first conducted through a cuvette, containing a plant, subsequently through the chemiluminescence analyzer. The gas was purified from NO by pulling it through charcoal column. The CO_2_/H2O gas exchange system GFS-3000 (Walz) was equipped with the LED-Array/PAM-Fluorometer 3056-FL for illumination and connected with an Arabidopsis Chamber 3010-A. Environmental parameters important for plant photosynthesis such as temperature, CO_2_ (400 ppm), relative humidity (50 %) and light. Temperature and light were dependent from the experimental setup. For the sunflecks experiment a plant was first adapted to ambient conditions (200 µmol photons m^-2^ s^-1^ PPFD and 22 °C) for 1 h afterwards a light stress was applied. A sunflecks pattern was created by increasing a light intensity to 1000 µmol photons m^-2^ s^-1^ PPFD and temperature to 30 °C for 10 min followed by returning both parameters back to ambient conditions for other 10 min. This pattern was repeated in total for four times. Additionally the emission of soil without a plant was measured and subtracted from plant emission.

### Chromatin immunoprecipitation sequencing (ChIP-seq)

#### Experimental design

Wild type and mutant plants were grown under chamber-controlled conditions (10/14 h light/dark) for 4 weeks. At this time plants achieved similar development stage. At midday (11 am: 5 h after turn on the light) plants were transferred either to dark (D, 0 µmol photons m^-2^ s^-1^ PPFD, 22 °C), low light (LL, 200 µmol photons m^-2^ s^-1^ PPFD, 22 °C) or to high light (HL, 1000 µmol photons m^-2^ s^-1^ PPFD, 30 °C) conditions for 4 h. After all, they were harvested and immediately cross-linked.

#### Cross-linking

1-2 g of Arabidopsis leaves were put in a 50 ml plastic tube and fill up with 30 ml precooled crosslinking buffer containing 1% formaldehyde. Concentration of suitable formaldehyde amount was obtained experimentally. The tubes were put in desiccator and vacuum was applied for 10 min. Crosslinking was stopped by adding to each tube glycine with the end concentration of 0.125 M followed by vacuum infiltration for another 5 min. After that leaves were washed twice with cooled water and dried on paper towels. Collected material was frozen in liquid nitrogen and stored at -80 °C.

#### Antibody coupling to magnetic beads

For each IP 20 µl of magnetic beads A were used. Beads of one biological replicate were washed together by pipetting up and down for 4 times with 1 ml buffer RIPA plus protease inhibitor. After, beads were suspended in the same volume with RIPA. Following antibodies were used to immunoprecipitate protein-DNA complex: anti-H3K9/14ac antibody (1 µg/IP), anti-H3K9ac antibody (1 µg/IP), IgG antibody (1 µg/IP, negative control) were added. Coupling of the antibodies to the beads was performed at 4°C on a rotation platform for approximately 7 h. In between chromatin isolation steps were performed. After coupling the AB-coated beads were washed with 500 µl RIPA for 3 times and resuspended with the same buffer. The beads were divided into new clean tubes (20 µl/IP).

#### Chromatin isolation

Leaves were ground to fine powder with mortar and pistil in liquid nitrogen. 2.3 g and 1.3 g of grounded material for ChIP-qPCR and ChIP-seq respectively were transferred in a 50 ml plastic tube and mixed with 20 ml Extraction buffer # 1. The suspension was incubated for 15-20 min at rotation platform at 4°C, followed by centrifugation at 4°C and 2800 g for 20 min. After that, supernatant was removed and pellet was suspended in total with 3 ml NRBT buffer. First 1 ml of buffer was added, pellet was suspended with a pipet tip, and then the rest 2 ml were added. Further, the nuclei were extracted using the same procedure as described before.

#### Sonication

After nuclei were isolated, they were carefully suspended (avoiding foam formation) with nuclei sonication buffer. Bioruptor® Pico ultrasonic bath and Covaris E220 Evolution were used to shear isolated chromatin for ChIP-qPCR and ChIP-seq, respectively. To perform DNA shearing for ChIP-qPCR 320 µl of sonication buffer were added to nuclei and transferred to 1.5 ml Bioruptor Microtubes (Cat No. C30010016). In total, 14 cycles with 30 sec ON/OFF was used. To perform DNA shearing for ChIP-seq nuclei were resuspended in 220 µl of sonication buffer and transferred to micro Tube AFA Fiber Pre-SlitSnap Cap (Cat No. 520245). Following sonication conditions were used: PIP - 175, DF – 10 %, CPB - 200, 600 sec. After this, sonicated samples were spun for 5 min at 16000 g and 4°C and the supernatant was used directly for immunoprecipitation assay or for the detection of shearing efficiency.

#### Shearing efficiency

50 µl and 20 µl of sonicated chromatin for ChIP-qCR and ChIP-seq, respectively, were diluted to 100 µl with sonication buffer. De-crosslinking was performed by adding 6 µl of 5 M NaCl and samples were incubated for 20 min at 95 °C and 1300 rpm. After that, 2 µl of RNaseA were added and samples were incubated for another 40 min at 37 °C and 1300 rpm. DNA was extracted using MinElute PCR purification kit (Qiagen, Cat No. 28004) or by phenol-chloroform followed by ethanol precipitation. DNA was eluted with 11 µl of dH2O. Concentration was measured using NanoDrop.

#### Immunoprecipitation and reverse crosslinking

For ChIP-qPCR 50 µl of sonicated chromatin were diluted with 200 µl buffer RIPA (1:5). 10 µl of diluted chromatin were saved as ‘‘Input’’ (4%). For ChIP-seq sonicated chromatin was diluted 1:10 with RIPA and 10 % were saved as “Input”. The diluted chromatin was added to AB-coated beads and incubated over night at 4 °C on a rotating platform. After, the beads were washed for 2 times with 1 ml of following buffers: low salt buffer, high salt buffer, LiCl buffer and TE buffer. Each wash step was performed on a rotating platform for 5 min at 4 °C. Immunoprecipitated chromatin (IP) was eluted with 125 µl of elution buffer plus proteinase inhibitor incubating at thermoblock for 15 min at 1200 rpm and 65 °C. Elution was performed twice and bough eluates were mixed together. For de-crosslinking to each ‘‘Input’’ sample elution buffer was added to reach the same volume as for IP samples (250 µl). De-crosslinking was performed by mixing each sample with 10 µl of 5 M NaCl (0.2 M NaCl end concentration) and incubating at 65 °C for at least 4-5 h and 1300 rpm. After that, samples were treated for 1 h with 4 µl of RNaseA (10 mg/ml) at 37 °C. Proteinase K treatment was performed for another two more hours by adding 2 µl Proteinase K (19.2 mg/ml), 5 µl of 0.5 M EDTA and 10 µl of 1 M Tris-HCl (pH 6.5). DNA was purified as described above. The DNA was eluted with 21 µl of dH2O for ChIP-qPCR or 15 µl of EB elution buffer (Qiagen, Cat No. 154035622) for ChIP-seq. DNA concentration was measured using Qubit(tm) dsDNA HS Assay Kit (Cat No. Q32851).

#### ChIP-seq

Size selection of fragmented DNA was additionally performed before library preparation using AMPure XP beads (Beckman Coulter, Cat No. A63881). 21 µl of magnetic beads (1.4:1, ratio of beads to sample) were added to each sample and incubated for 10 min at RT. After, beads were placed to a magnetic stand and the supernatant was disposed. Beads were washed three times with 20 µl of 80 % ethanol and dried. DNA was eluted with 12 µl of EB elution buffer by incubation the beads for 3 min. The size of immunoprecipitated and “Input” samples was analyzed using Agilent High Sensitivity DNA Kit (Cat No. 5067-4626) at Agilent 2100 Bioanalyzer according to the manufacturing instructions. Library preparation and deep sequencing was performed by IGA Technology Services (https://igatechnology.com/) using NextSeq500 and 30 M (75 bp) reads.

#### ChIP-seq data analysis

The ChIP-seq reads were aligned against TAIR10 reference genome assembly for Arabidopsis thaliana (accessed on May 14th 2018) using bowtie2-2.3.4.1 (Langmead et al., 2009). After quality-based filtering with samtools-1.8 (Li et al., 2009) using -q 2, MACS-1.4.2 (Zhang et al., 2008) was applied for peak calling against the input controls (whole DNA, no ChIP), with genome size 1.35e8, model fold 8,100, fragment size 150 and p-value cutoff 1e-5. Differential analysis between groups was performed based on the DESeq2 method (Love et al., 2014) using DiffBind 2.12.0 (Ross-Innes et al., 2012; Stark and Brown, 2019), re-centering the peaks at summits and setting the width of consensus peaks to the maximum fragment size estimate by MACS-1.4.2. The alignment format conversion required for DiffBind was done with samtools-1.8 (Li et al., 2009). Differential peaks with adjusted p-value (false discovery rate method, FDR) < 0.05 were used for further analyses. Venn diagrams were made with the R package limma, version 3.40.2 (Ritchie et al., 2015), significance of overlaps was assessed with fisher.test in R version 3.6.0 (Team, 2019). Principal component analysis by prcomp and plot functions were employed in R version 3.6.0 (Team, 2019) for visualization of the normalized count data from DiffBind. Read counts for specific genomic locations were queried by samtools-1.8 (Li et al., 2009) and scaled to a common library size of 10e6 for co-visualization of ChIP-seq and RNA-seq output.

#### Functional enrichment analysis

Gene Ontology (GO) term enrichment was computed in R version 3.6.0 (Team, 2019), applying fisher.test and p.adjust with FDR. The GO terms and annotated genes were taken from org.At.tairGO2ALLTAIRS in the org.At.tair.db R package, version 3.8.2 (Carlson, 2019b). The description of the GO term was obtained from the GO.db R package version 3.8.2 (Carlson, 2019a). Significantly enriched GO terms (FDR < 0.05) were subjected to multi-dimensional scaling (MDS) analysis by cmdscale in R version 3.6.0 (Team, 2019) using the function dist with method “binary” on their profiles of differential genes. Significantly enriched GO terms from the biological process ontology were plotted with respect to the first two MDS coordinates and colored according to their ancestors among the top level biological process terms, which were classified into five broader categories (response to stimulus: GO:0002376, GO:0023052, GO:0050896; localization: GO:0051179; growth and development: GO:0000003, GO:0008283, GO:0022414, GO:0032501, GO:0032502, GO:0040007, GO:0071840; metabolic process: GO:0008152; other: GO:0001906, GO:0006791, GO:0006794, GO:0007610, GO:0009758, GO:0009987, GO:0015976, GO:0019740, GO:0022610, GO:0040011, GO:0043473, GO:0044848, GO:0048511, GO:0051704, GO:0065007 GO:0098743, GO:0098754, GO:0110148). The visualization was achieved by the R package ggplot2, version 3.1.1 (Wickham, 2016), and scatterpie, version 0.1.4 (Yu, 2019), as well as the barplot function of R version 3.6.0 (Team, 2019).

#### RNA-seq

Sequencing libraries were generated from poly(A)-enriched RNA using the NEBNext Ultra II Directional RNA Library Prep kit (New England Biolabs) according to the manufacturer’s instructions, and sequenced on an HiSeqV4 instrument (Illumina) as 100bp single-end reads in a 24-plex pool. Reads were mapped to the TAIR10 reference of *Arabidopsis thaliana* annotated genes (www.arabidopsis.org) using STAR (v2.5.2a) (Dobin et al., 2013). Read quantifications were generated using kallisto (v0.43.1) (Bray et al. 2016). Differential expression analysis was performed using the DESeq2 package (v1.18.1) with default settings (Love et al., 2014) in R (v3.4.4) (Team, 2017). Genes were considered as differentially expressed if the expression level between samples differed by more than 2-fold and if the Benjamini-Hochberg-adjusted *p*-value was < 0.1.

#### Data availability

ChIP-seq and RNA-seq data will be available in the ArrayExpress functional genomics database.

## Materials Availability statement

All unique/stable reagents/plant material generated in this study are available from the Corresponding Author without restriction.

## Data and Code Availability Statements

The ChIP-seq and RNA-seq data generated during this study will be available at ArrayExpress functional genomics database.

## Supplemental Information

**Supplementary Table S1:** ChIP-seq and RNA-seq sample and alignment information.

**Supplementary Table S2:** Differential genes of LL vs. D and mutant vs. wild type comparisons for ChIP-seq and RNA-seq data. Up-regulation of gene expression or acetylation peaks close to transcription start sites in the first vs. the second condition is indicated by 1, down-regulation by -1 (FDR-adjusted p-value < 0.05). Genes with both up- and down-regulated acetylation peaks are marked by -1/1.

**Supplementary Table S3:** GO term enrichment of LL vs. D comparisons for ChIP-seq data. For each GO term identifier, the table gives raw and FDR-adjusted enrichment p-values, the descriptive name and the identifiers of genes from this term that are present in the list of differential genes. There is a separate sheet for enrichment analyses of each of the following LL vs. D differential gene lists derived from ChIP-seq data: up in all three genotypes (wt, *gsnor, hda6*), down in all three genotypes, up in both mutants (*gsnor, hda6*) but not wt, down in both mutants but not wt, up in wt but not any mutant, down in wt but not any mutant.

**Supplementary Table S4:** GO term enrichment of mutant vs. wild type comparisons for ChIP-seq data. This table shows enrichment analyses (performed as in Supplementary Table S3) for the following mutant (*gsnor, hda6*) vs. wt differential gene lists in LL condition derived from ChIP-seq data: up in both mutants, down in both mutants.

**Supplementary Table S5:** GO term enrichment of LL vs. D comparisons for RNA-seq data. This table contains the RNA-seq results for an analysis equivalent to Supplementary Table S3.

**Supplementary Table S6:** GO term enrichment of mutant vs. wild type comparisons for RNA-seq data. This table contains the RNA-seq results for an analysis equivalent to Supplementary Table S4.

## Supplement

**Supplementary Figure 1:**
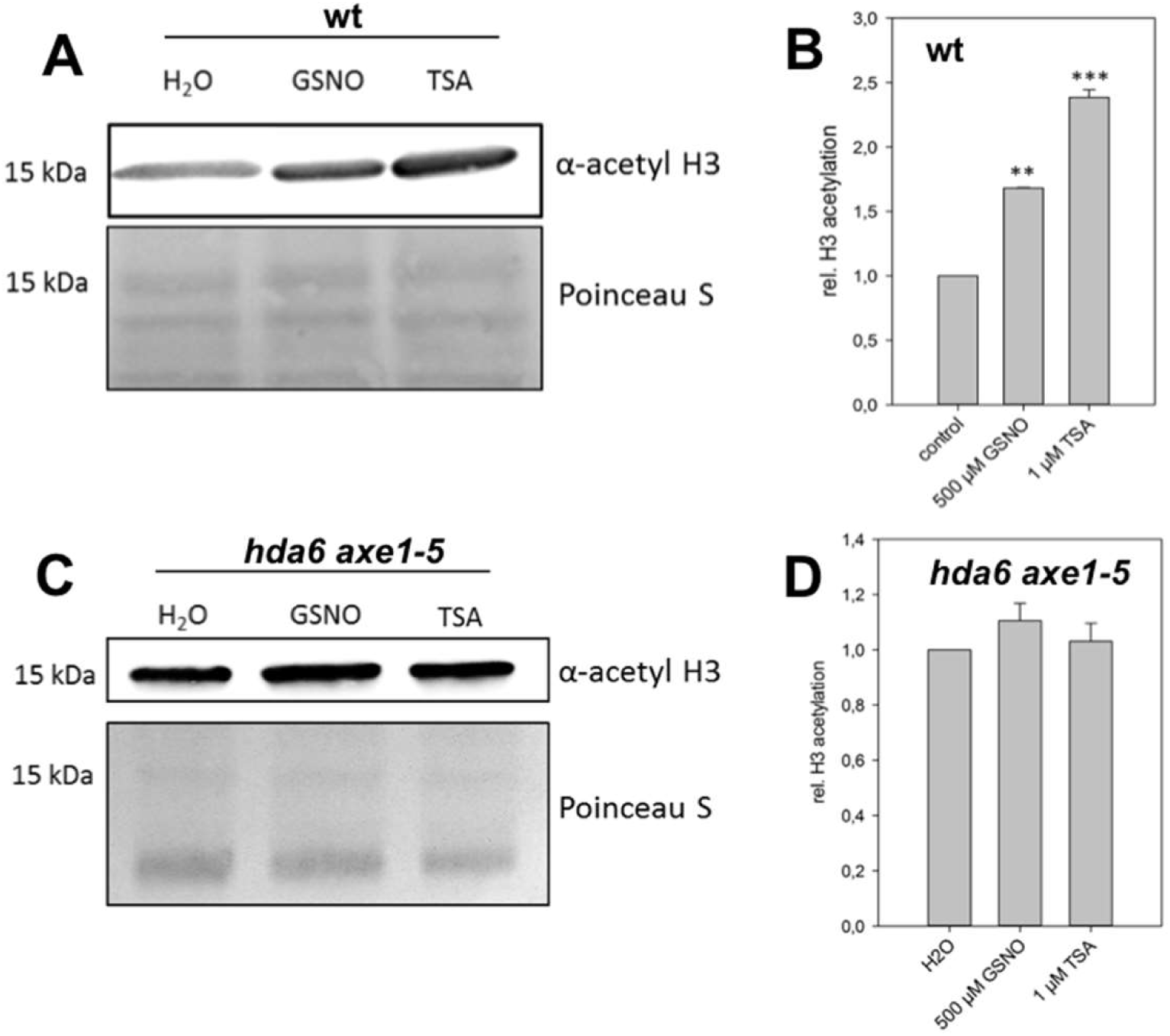
Comparison of H3 acetylation in wt and *hda6* suspension cells after GSNO treatment. A and C) Western-Blot analysis of GSNO- and TSA-treated wt and *hda6* cells. Nuclear extracts were separated by SDS-PAGE and blotted. The membrane was probed with an anti-acetyl H3 primary antibody and a secondary antibody coupled to HRP. Shown is one representative experiment. B and D) Quantification of A and C. Signal intensity was determined with Image J software. Shown is the mean ± SEM of three experiments. **P < 0.01, ***P < 0.001, student’s t-test. These experiments were done by Alexandra Ageeva under my supervision.

**Supplementary Figure 2:**
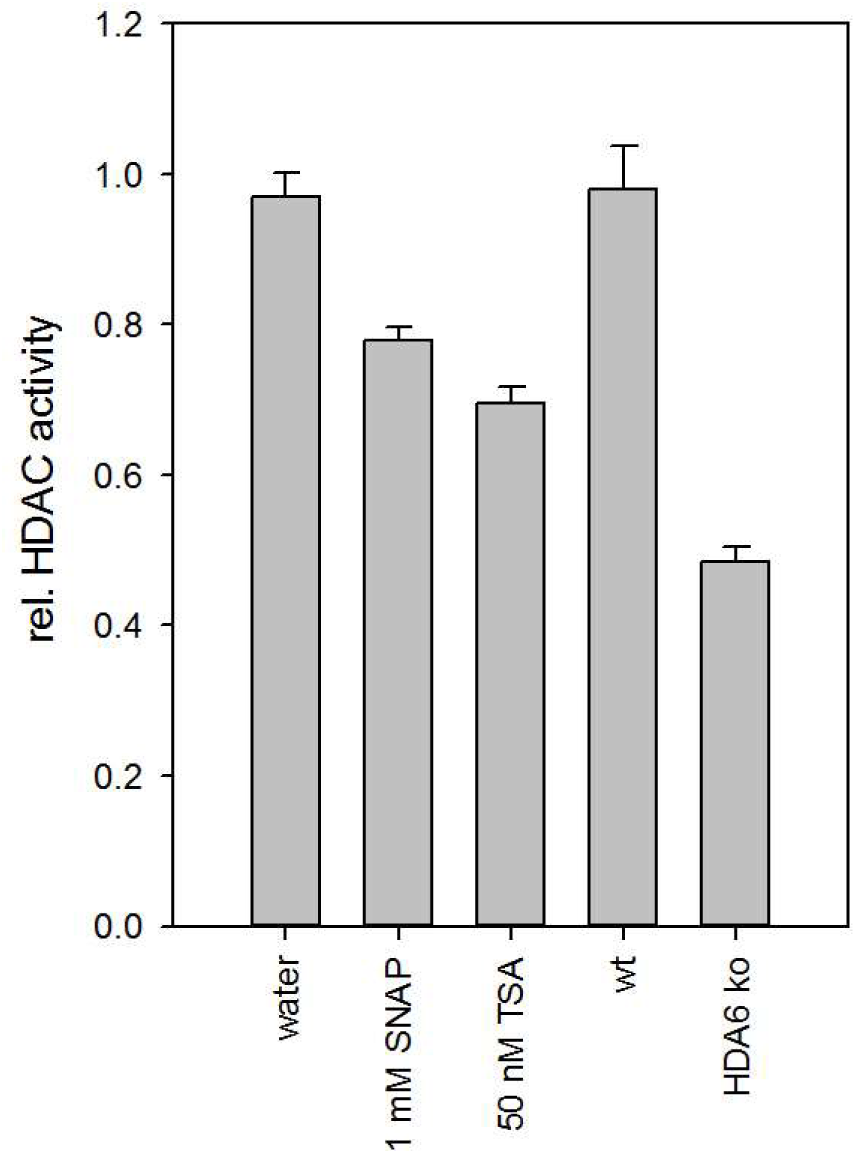
HDAC activity in nuclear extracts of wt and *hda6* cell culture. Nuclear extracts were prepared according to section 5.4.2 and HDAC activity was measured as described. Values are normalized to water treatment or wt. Shown is the mean of two independent experiments with three technical replicates each. *P-value < 0.05, **P-value < 0.01.

**Supplementary Figure 3:**
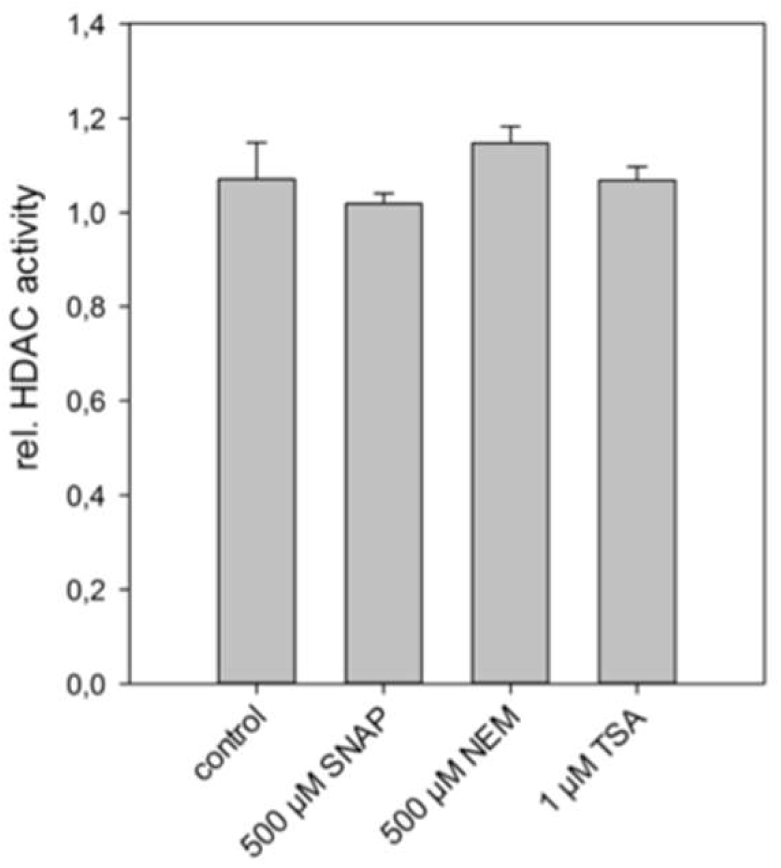
Insensitivity of HDAC activity in *hda6* suspension cells towards cysteine modifications and TSA. Nuclear extracts from *hda6* suspension cells were incubated with 500 µM SNAP, 500 µM NEM and 1 µM TSA and HDAC activity was measured over 90 min. Values are normalized to control treatment (water). Shown is the mean ± SEM of three independent preparations of nuclear extract.

